# Genetic Dissection of Grain Yield and Correlated Proxy Traits Under Suboptimal Conditions

**DOI:** 10.64898/2026.04.22.720082

**Authors:** Yan-Cheng Lin, Claude Urbany, Antonina Shlykova, Armin Hölker, Milena Ouzunova, Thomas Presterl, Torsten Pook, Manfred Mayer, Sebastian Urzinger, Chris-Carolin Schön

## Abstract

Securing sustainable crop production requires the genetic improvement of abiotic stress tolerance. Due to the broad range of environmental factors causing abiotic stress and complex genotype-by-environment interactions, it is crucial to understand the genetic basis of crop yield under suboptimal conditions. Here, we developed a dent maize Multi-parent Advanced Generation Inter-Cross (MAGIC) population comprising 388 doubled haploid (DH) lines. The population was derived from eight founders with varying stress tolerance, selected from a dent diversity panel evaluated for yield performance across a wide range of European environments. The MAGIC DH lines were genotyped via whole-genome sequencing (∼5X coverage) and evaluated in seven testcross and 14 line *per se* trials, for grain dry matter yield, leaf senescence, leaf rolling, anthesis-silking interval, and six additional agronomic traits. Genetic dissection identified 22 grain yield QTL, explaining 45% of the genetic variance. Under heat and drought stress, testcross grain yield correlated significantly with leaf senescence and leaf rolling measured in line *per se* trials. Bivariate multi-trait analysis showed that alleles for delayed senescence and reduced rolling at detected QTL generally exhibited positive effects on grain yield, suggesting that accumulating these favorable alleles could enhance yield performance. Incorporating these proxies into multi-trait genomic prediction models improved yield prediction accuracy, although gains were constrained by modest trait correlations. Given the comprehensive data, we also provide recommendations for optimizing sequencing depth and QTL mapping strategies in experimental maize populations.

**Key message:** This eight-founder MAGIC population represents a powerful resource for dissecting complex traits in maize, assessing the utility of drought proxy traits, and optimizing low-coverage whole-genome sequencing approaches.

## Introduction

Breeding for tolerance to water-deficit conditions has become a critical strategy to enhance resilience of our crops to climate change and ensure food security. However, directly selecting for grain yield under suboptimal conditions has remained difficult due to high variability of potential environmental scenarios and complex genotype-by-environment (G×E) interactions affecting yield performance. Indirect selection for grain yield either by genome-based strategies or facilitated by the evaluation of proxy traits that can be measured at high-throughput might attenuate these limitations.

During the vegetative stage, leaf rolling has been suggested as a physiological indicator of drought response in maize (Bänziger et al. 2000), but its relevance for selecting drought-tolerant genotypes has remained elusive. Saglam et al. (2014) compared two cultivars with different drought responses and reported that leaf rolling reduces water loss and mitigates stress damage by limiting transpiration in the tolerant genotype. In contrast, Effendi et al. (2019) showed that drought-tolerant varieties exhibited less leaf rolling, suggesting that excessive rolling could signal susceptibility rather than tolerance. During the reproductive stage, asynchronous development of anthesis and silking under stress has been shown to lead to severe yield loss (Bruce et al. 2002). However, modern maize hybrids have been selected for short ASI (Araus et al. 2012), making it less relevant when selecting for yield improvement under challenging conditions. During the grain-filling period, delayed senescence (stay-green) has been suggested as one of the mechanisms underlying stress tolerance (Tollenaar and Wu 1999). Genotypes with longer stay-green have been shown to have higher grain yield under drought-heat conditions (Cerrudo et al. 2017). However, the stay-green phenotype may hinder grain dry-down when green leaf area is retained beyond maturity (Lee and Tollenaar 2007).

In a seminal study aimed at understanding the genetic control of grain yield in heat-and drought-prone environments, a diversity panel (DROPS panel) of 244 dent maize hybrids was evaluated under irrigated and rainfed conditions across nine European sites in the years 2012 and 2013 and one site in Chile in 2013, totaling 29 field experiments (Millet et al. 2016). The extensive multi-environment dataset facilitated the characterization of QTL allelic effects on yield in relation to environmental variables and supported the development of genomic prediction models that incorporated genotype-by-environment interactions (Millet et al. 2019). Building on the results of this study, we developed a Multi-parent Advanced Generation Inter-Cross (MAGIC) population using eight inbred lines selected from the diversity panel for their contrasting performance in drought and heat prone environments.

We chose the MAGIC design due to its favorable properties for QTL mapping and genomic prediction (Scott et al. 2020). MAGIC populations are created by inter-mating several inbred founders over multiple generations before deriving homozygous inbred lines for phenotyping. This inter-mating process leads to a population with genomes that form fine-scale mosaics, reflecting balanced contributions from all founder lines (Huang et al. 2015). Compared to diversity panels, MAGIC populations have minimal population structure. Known founder genotypes and balanced allele frequencies facilitate accurate imputation when using whole-genome sequencing as a genotyping platform. Compared to bi-parental populations, MAGIC populations exhibit significantly higher genomic diversity and recombination.

In maize, several MAGIC populations have been developed for genetic studies. They offered high power and resolution for QTL mapping and facilitated the identification of candidate genes (Dell’Acqua et al. 2015; Ferguson et al. 2025). The genetic control of early-stage drought tolerance (Rida et al. 2021) and leaf senescence (Caicedo et al. 2021) has also been studied using MAGIC populations. Genomic prediction in MAGIC populations has been shown to have high accuracy for important agronomic traits (Michel et al. 2022). Moreover, Hudson et al. (2022) used a MAGIC population to study genotype-by-environment (G×E) interactions.

Here, we evaluated an eight-founder MAGIC population across 21 diverse field experiments. Our objectives were to: (i) assess the efficiency of shallow-seq as a genotyping platform and suggest best practices, (ii) characterize the genetic and phenotypic variation of the developed MAGIC population for important agronomic traits including grain yield under suboptimal conditions, and (iii) evaluate the relevance of proxy traits for grain yield improvement.

## Materials and methods

### Genetic material

A MAGIC population comprising 388 doubled-haploid (DH) lines was developed by intercrossing eight dent maize inbred lines using three funnel crossing schemes (subfamilies), as detailed in Table S1. The eight founder lines, B106, B107, F888, FC1890, Lo1056, Lo1270, Lo1290, and PHG83, were selected from the DROPS panel based on their grain yield performance in ten testcross trials conducted by Millet et al. (2016). These founders exhibited similar grain yield in irrigated trials, while showing variable performance in rainfed trials (Fig. S1A). Founders FC1890, B107, and PHG83 belong to the Iodent heterotic group, Lo1056, Lo1270, and Lo1290 belong to the Lancaster group, and F888 and B106 have been categorized as other Non-Stiff Stalk (NSS) material (Fig. S1B). The eight lines were selected for moderate variation in flowering time, with a maximum difference of about eight days under irrigation and 11 days under rainfed conditions (Millet et al. 2016).

### Field experiments

The 388 MAGIC DH lines, their founder lines, together with 18 elite inbred lines proprietary to KWS SAAT SE were phenotyped in 21 field trials conducted between 2020 and 2023 in seven locations in Germany (4), Hungary (2), and Italy (1). Seven trials assessed testcross performance using the flint inbred line UH007 as the tester, consistent with the DROPS project, while fourteen trials evaluated line *per se* performance. All trials were arranged in generalized alpha-lattice designs with two replications and block sizes of either five or ten plots. In addition to the final set of 388 MAGIC DH lines, trials included DH lines that did not pass marker-based identity filtering and were treated as checks in subsequent phenotypic analyses.

Two plot types were used: observation plots (3 m long, 0.75 m row spacing, planting density of 8.8 plants m^-2^) and yield plots (6 m long, 0.75 m row spacing, 11 plants m^-2^). All yield plots were planted as double-rows. Observation plots were single-row, except for two trials with double-rows in 2023. Sowing, fertilization, and plant protection followed standard agricultural practices at the respective experimental stations. Some trials were conducted as rainfed experiments on sandy soils to simulate water-limited conditions. Summary information on all field experiments is provided in Table S2.

Weather data were obtained for each trial from DTN ClearAg™ (https://www.dtn.com/agriculture/agribusiness/clearag). Data covered a 60-day window spanning 30 days before and 30 days after male flowering (median across all plots) and are provided in File S1. To specify the environmental conditions associated with water and temperature stress during this period, we summarized precipitation, reference evapotranspiration, (ET_0_, calculated using the FAO-56 Penman-Monteith equation), and maximum temperature in Fig. S2.

### Trait assessment

Ten traits were measured (Table S2), including grain dry matter yield (GDY, calibrated to 85% dry matter content, t/ha), grain dry matter content (GDC, %), early plant height (PH_V6, measured at the V6 stage from the soil surface to the tip of the tallest leaf when stretched, averaged over three plants per plot, cm), final plant height (PH_final, measured from the soil surface to the ligule of the flag leaf, averaged over three plants per plot, cm), ear height (EH, measured from the soil surface to the ligule of the upper cob, averaged over three plants per plot, cm), male flowering (MF, number of days from sowing to 50% of plants shedding pollen in a plot, days) and female flowering (FF, number of days from sowing to 50% of plants silking in a plot, days).

Proxy traits included a leaf senescence score (LS, scored 4-6 weeks after flowering on a 1-9 scale, where 1 = very green and 9 = complete senescence), a leaf rolling score (LR, scored on a hot dry day on a 1-9 scale, where 1 = no leaf rolling and 9 = severe rolling across all plants), and the anthesis-silking interval (ASI, days). Leaf senescence and leaf rolling were recorded only when phenotypic variation was visible.

### Phenotypic data analysis

The statistical model for analyzing the phenotypic data is given by:

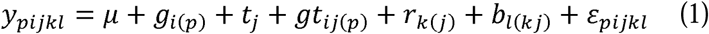

Here, *y*_*pijkl*_ is the phenotypic plot-level observation; 𝜇 denotes the overall mean; *g*_*i(p)*_ is the effect of genotype 𝑖 nested in group *p*, where *p* = 1 for MAGIC DH lines and *p* = 2 for founders and checks. For variance component estimation, *g*_*i(p)*_ was treated as random effect for *p* = 1 and as fixed for *p* = 2. For obtaining adjusted entry means across trials, *g*_*i(p)*_ was treated as fixed for all genotypes. 𝑡_𝑗_ is the random effect of trial 𝑗; 𝑔𝑡_𝑖𝑗(*p*)_ is the random interaction effect of genotype 𝑖 and trial 𝑗 with heterogeneous variances 𝜎^2^ among groups; 𝑟_𝑘(𝑗)_ and 𝑏_𝑙(𝑘𝑗)_ are random effects of replication 𝑘 nested in trial 𝑗 and incomplete block 𝑙 nested in replication 𝑘 in trial 𝑗; 𝜀_*pijkl*_ is the random residual error of observation *p*𝑖𝑗𝑘𝑙, with 𝜀_*pijkl*_ ∼ iid 𝑁(0, 𝜎^2^). ASReml (v4.2) (Butler et al. 2017) was used to fit linear mixed models, estimate variance components and compute adjusted entry means.

Adjusted entry means for individual trials were calculated by excluding trial (𝑡_𝑗_) and genotype-by-trial (𝑔𝑡_𝑖𝑗(*p*)_) effects from Eq. (1). Trait heritabilities and their 95% confidence intervals were estimated on an entry-mean basis following (Hallauer et al. 2010) and Knapp et al. (1985), respectively. Phenotypic correlations were calculated from adjusted entry means of individual trials using Pearson correlation coefficients with false discovery rate (FDR) correction (Benjamini and Hochberg 1995) accounting for multiple testing.

### Whole genome short-read sequencing

The eight founders of the MAGIC population were sequenced at high depth (>50X), while the 388 MAGIC DH lines were sequenced at an average genome sequencing depth of 5X (shallow-seq) using 150-bp paired-end short reads generated with the Illumina NovaSeq 6000 platform. For quality assessment, the eight founder lines and a subset of 64 DH lines were also genotyped using the 600k Affymetrix Axiom Maize Array (Unterseer et al. 2014).

Initially, variant calling was performed using whole genome sequences of the eight founder lines. The raw reads were preprocessed by removing PCR duplicates using hts_SuperDeduper (v1.3.0) (Bonfield et al. 2021), trimming adapters and low-quality sequences with Trimmomatic (v0.39) (Bolger et al. 2014), and correcting base errors with Lighter (v1.1.2) (Song et al. 2014). Reads were aligned to the B73v5 reference genome (Hufford et al. 2021) using bwa (v0.7.17) (Li and Durbin 2009). Alignment (BAM) files were sorted using SAMtools (v1.9) (Li et al. 2009) and duplicate reads were marked with picard MarkDuplicates (v2.26.0) (https://broadinstitute.github.io/picard/). Variant calling was performed with freebayes (v1.3.5) (Garrison and Marth 2012). Variants located in low-complexity regions annotated by sdust (Morgulis et al. 2006) were removed. Bi-allelic homozygous SNPs with complete data across all founders (8.3 M) were selected as reference for the downstream variant calling in the DH lines.

Reads of DH lines were preprocessed with the same pipeline for trimming, base-correction, duplicate removal and alignment. Variant calling was performed with freebayes, using the founder-derived SNP positions as reference. Reads from founder lines were jointly called as additional samples. SNP loci were excluded if founders showed inconsistent genotypes between the two variant-calling runs, heterozygosity was greater than 10%, or missing rates exceeded 50% in the MAGIC population. Further filtering removed SNPs located in annotated transposable elements (*Zm-B73-REFERENCE-NAM-5.0.TE.gff3.gz*, downloaded from https://download.maizegdb.org/). The final dataset was imputed using the R package HBimpute (Pook et al. 2021) and SNPs with minor allele frequency (MAF) < 0.05 were discarded, yielding 2,717,240 high-quality SNPs for downstream analyses.

To assess the efficiency of shallow-seq, we down-sampled reads from 31 DH lines with high depth (>8X) to depths of 0.1X, 0.5X, 1X, 2X, 4X, and 8X using Seqkit (Shen et al. 2016). At each depth, variant calling followed the same reference SNP guided pipeline described above. Imputation was conducted on all SNP datasets without prior SNP filtering using Beagle 5.4 (Browning et al. 2018). For each sequencing depth, we evaluated four metrics: (1) genome coverage, defined as the proportion of the genome covered by reads, (2) genotyping rate, defined as the proportion of SNPs genotyped per individual, (3) number of usable SNPs, defined as SNPs with a missing rate below 50%, and (4) genotyping error rate, calculated as the concordance with the 600k SNP array genotypes. Summary statistics were computed using SAMtools (Li et al. 2009) and VCFtools (Danecek et al. 2011).

### Population structure and LD decay

Population structure of the DH lines was assessed using a subset of 20,974 SNPs selected from the full SNP set (2.7 M) via LD pruning (*r^2^*< 0.4 within a 500 kb window) using PLINK (Purcell et al. 2007), balancing genome coverage and minimal LD. Pairwise genetic distances between individuals were computed using modified Rogers distances (Wright 1978). Principal Coordinate Analysis (PCoA) was conducted using the *pcoa()* function in R. A neighbor-joining phylogenetic tree (Saitou and Nei 1987) was constructed utilizing the *nj()* function from package ape (Paradis and Schliep 2019) and visualized with the phytools (Revell 2012) package.

For LD decay analysis, the full SNP dataset was thinned by selecting every 10th SNP to reduce computational demands. Pairwise LD (*r^2^*) between SNPs was computed using PopLDdecay (Zhang et al. 2019). LD decay distance was estimated for an *r^2^* threshold of 0.2 (Hill and Weir 1988).

### Genomic mosaic reconstruction and recombination estimation

Founder haplotype (FH) probability, *i.e.*, the likelihood that a specific allele in a progeny line originated from a given founder, was estimated using the R-package HaploBlocker (v1.7.1) (Pook et al. 2019) with function *founder_detection()*, which employs a Hidden Markov Model (HMM), similar to the approach used in mpmap2 (Shah et al. 2019).

HaploBlocker improves upon mpmap2 by using a haplotype library constructed from both founders and progeny, which adjusts emission probabilities when shared haplotype blocks are detected. To reduce computational load and minimize noise from genotyping errors, the 2.7 million SNP dataset was partitioned into 50 subsets by assigning every 50th marker to the same subset. Founder probabilities were computed independently for each subset and subsequently averaged across a 50-marker sliding window. While this approach entails a reduction in mapping resolution, it significantly mitigates the influence of consecutive genotyping errors.

Founders may share chromosome segments that are identical by descent (IBD). Pairwise IBD regions of founders were detected using *phasedibd* (Freyman et al. 2021) with default settings, and IBD proportions were estimated based on the length of IBD regions on the linkage map. As these segments likely reflect common ancestry, founder alleles were merged into an ancestral allele if the corresponding chromosome regions were IBD between founders. For each DH line, ancestral haplotype (AH) probabilities were obtained by summing founder haplotype probabilities within each ancestral group.

Recombination event counts were estimated using the haploRILs R package (https://github.com/GoliczGenomeLab/haploRILs). The method compares the genomic composition of each DH line to the founder genomes by assigning founder haplotype blocks along each chromosome. The SNP dataset was divided into 50 subsets (every 50th marker), and estimations were performed independently for each subset using the settings *nSnp = 20*, *step = 1*, and *k = 1*. Final recombination counts per chromosome per individual were calculated as the average across all 50 runs.

### Genome-wide association (GWA) analysis in individual trials

We first performed a GWA analysis on 388 DH lines using bi-allelic SNPs. The dataset was pruned with PLINK (Purcell et al. 2007) to remove redundant markers (*r^2^* > 0.99 within a 300 kb window). Given the high marker density, SNPs that showed *r^2^* < 0.8 with all other markers within 10 Mb distance were considered erroneous and excluded. This resulted in a final set of 359,141 SNPs. Association analyses were conducted on the adjusted entry means from individual trials and means across trials using vcf2gwas (Vogt et al. 2022), which implements a mixed linear model of GEMMA (Zhou and Stephens 2012):

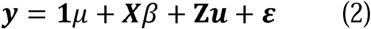

where 𝒚 is the 𝑛-dimensional vector of adjusted entry means, with 𝑛 being the number of MAGIC lines (𝑛 = 388); 𝟏 is a vector of ones; 𝜇 is the overall mean; 𝛽 is the fixed effect of the tested SNP marker; 𝑿 is the vector of corresponding SNP genotype scores coded as 0 and 2; 𝒖 is the 𝑛-dimensional vector of random polygenic effects, where 𝒖∼𝑁(𝟎, 𝐊𝜎_g_^2^), with 𝜎_g_^2^ being the genetic variance pertaining to the model. 𝐊 denotes the 𝑛 × 𝑛 realized genomic relationship matrix, computed using the method of Astle and Balding (2009) implemented with the leave-one-chromosome-out (LOCO) approach described by Yang et al. (2014); 𝐙 is the design matrix for the polygenic effects in 𝒖; and 𝜺 is the 𝑛-dimensional vector of random residual effects, where 𝜺∼𝑁(𝟎, 𝐈_𝐧_𝜎^𝟐^) with 𝐈_n_ being an 𝑛 × 𝑛 identity matrix and 𝜎^2^ the residual variance.

SNP-trait associations were assessed using the Wald test (Wald 1943), and significance was determined at a 1% false discovery rate (FDR) (Benjamini and Hochberg 1995). Significant SNPs with *r^2^* > 0.2 within 3 Mb distance were merged into a single QTL, with the SNP showing the lowest *p*-value designated as the lead SNP. All lead SNPs were fitted in a multi-locus model using stepwise backward elimination, with each SNP tested as last fixed term using the Wald test. SNPs with *p*-values ≥ 0.01 were iteratively removed until all remaining SNPs met the threshold, resulting in a specific QTL set (𝑸) for each trait in each trial.

QTL heritability, i.e., the proportion of phenotypic variance explained by individual QTL, was estimated by treating QTL effects as random and using centered and scaled genotype coding to calculate their variance relative to the total phenotypic variance, following Wang et al. (2022) with the following model:

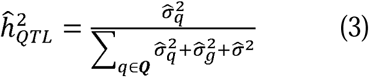

where 𝜎^^2^ is the variance estimate of a particular QTL; 𝑸 denotes the QTL set of a trait in a trial; 𝜎^^2^ and 𝜎^^2^ are the variance estimates of polygenic effects and residuals, respectively.

### Multi-trial model for final QTL set

To consolidate QTL for each trait, we performed stepwise backward elimination on all QTL detected from individual trial analyses, adapting the multi-locus, multi-trial model from Millet et al. (2016):

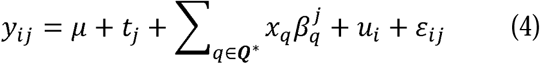

Where *y*_𝑖𝑗_is the adjusted mean of genotype 𝑖 in trial 𝑗; 𝜇 is the overall mean; 𝑡_𝑗_is the fixed effect of trial 𝑗; 𝑥_𝑞_ is the SNP genotype score (0 or 2); 𝛽*_q_*^𝑗^ is the fixed effect of SNP 𝑞 in trial 𝑗; 𝑢_𝑖_ is the random polygenic effect of genotype 𝑖. The kinship matrix 𝐊^∗^ was calculated based on all chromosomes as multiple QTL are fitted in the model; 𝜀_𝑖𝑗_is the random residual error with trial-specific error variance. 𝑸^∗^ denotes the final QTL set.

To avoid collinearity, nearby QTL (*r^2^* > 0.8 and < 3 Mb) from different trials were grouped retaining only the marker with the lowest *p*-value for backward elimination. Stepwise backward elimination was then performed as described above to obtain the final QTL set (𝑸^∗^). The allelic effects of the QTL remaining in the model were decomposed into a main effect and a QTL-by-trial (QxT) interaction (𝛽*_q_*^𝑗^ = α_𝑞_ + α*_q_*^𝑗^ ), and Wald tests of α*_q_*^𝑗^ were used to identify consistent or trial-specific QTL (*p* < 0.01).

Given the heterogeneous residual variances in the multi-trial analyses, we estimated the proportion of total genetic variance captured by all QTL in the final set. QTL effects were treated as fixed, and the proportion was calculated based on the reduction in genetic variance (𝜎^^2^) comparing models with and without QTL. QTL confidence intervals were defined by the two most distant SNP markers in strong LD with the lead SNP (*r^2^* > 0.8). Allelic effects at the lead SNPs were obtained as best linear unbiased estimates (BLUEs) of 𝛽 in Eq. (2) in individual trials and Eq. (4) for the final QTL set across trials. A flowchart of the full QTL analysis is available in Fig. S3.

### QTL nomenclature

QTL nomenclature was adapted from the convention proposed by McCouch (1997); for example, *qGDY(LP)01A*, where “GDY” refers to the trait (grain dry matter yield), “LP” in parentheses indicates the line *per se* trial (TC for testcross), “01” indicates chromosome 1, “A” denotes the first QTL on that chromosome based on its physical position.

### IBD-based QTL mapping

Odell et al. (2022) highlighted the value of multiple mapping approaches for QTL detection in a MAGIC population. Therefore, we also used founder (FH) and ancestral haplotypes (AH) as genomic predictors for QTL analyses. For individual trials, QTL detection was performed using an IBD-based mixed linear model implemented in the R package statgenMPP (Li et al. 2022), in which the effects of founder or ancestral alleles at a putative QTL were modeled as random effects for hypothesis testing. On the multi-trial analysis, a backward QTL elimination procedure was applied, adapting the multi-allelic, multi-locus hypothesis testing framework of Li et al. (2024). Detailed methodological procedures are described in Method S1.

### Multi-trait GWA analysis

We studied potential pleiotropy of QTL for proxy traits and grain yield using bivariate multi-trait models. Each model included adjusted means of grain yield estimated across testcross trials together with adjusted means of one proxy trait estimated across line *per se* trials. The bivariate multi-trait model of Korte et al. (2012), was implemented using the R package StatgenQTLxT (https://github.com/Biometris/statgenQTLxT). Extending the model in Eq. (2), 𝒚 is an 2𝑛 𝑥 1 vector (𝑛 = 388) stacking the centered and scaled phenotypes of grain yield and the proxy trait; 𝜷 is a 2 𝑥 1 vector of fixed SNP effects; 𝑿 is a 2𝑛 𝑥 2 design matrix for fixed effects, constructed as 𝑰_2_ ⊗ 𝒙, where 𝒙 is the 𝑛 𝑥 1 vector of SNP genotype scores (coded as 0 and 2); 𝒖 is the 2𝑛 𝑥 1 vector of random polygenic effects, with 𝒖∼𝑁(𝟎, 𝚺_𝒈_ ⊗ 𝐊), where 𝚺_𝒈_ is a 2 𝑥 2 genetic variance-covariance matrix between traits and 𝐊 is the same 𝑛 𝑥 𝑛 kinship matrix used in Eq. (2), and; 𝐙 is the 𝑰_2𝑛_ design matrix, for the polygenic effects 𝒖; and 𝜺 is the 2𝑛 𝑥 1 vector of residuals, with 𝜺∼𝑁(𝟎, 𝚺_𝜺_ ⊗ 𝑰_𝑛_), where 𝚺_𝜀_ is the 2 𝑥 2 residual variance-covariance matrix between traits. Both 𝚺_𝒈_and 𝚺_𝜺_ were modeled as unstructured matrices.

### IBS and IBD-based genomic prediction

Genomic best linear unbiased prediction (GBLUP) was used to obtain the genetic values of the 388 DH lines. The model was a modification of Eq. (2) excluding the fixed marker effect. The kinship matrix 𝐊 was replaced by a SNP-based genomic relationship matrix (GRM), denoted 𝐔_𝐒_, calculated following method 1 described by VanRaden (2008). The GBLUP model was implemented using ASReml (v4.2) (Butler et al. 2017).

In addition to the SNP-based GRM, we constructed GRMs 𝐔_𝐅𝐇_and 𝐔_𝐀𝐇_using founder and ancestral haplotype probabilities. At each marker locus, each DH line was assigned a vector indicating the probability of carrying a specific founder or ancestral allele. The probability that two DH lines shared the same founder allele at a locus was calculated as the dot product of their probability vectors. The GRM was then calculated as 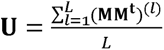, where 𝐌 is the 𝑛 𝑥 *p* probability matrix (𝑛 = number of DH lines, *p* = number of haplotypes) at a marker locus, and the GRM is average across all 𝐿 marker loci. For constructing the three GRMs we used the same evenly spaced 20K marker set described in Method S1. To test if the three GRMs differed significantly we performed a Mantel test (Mantel 1967).

### Multi-trait genomic prediction

A multivariate linear mixed model was implemented in ASReml (v4.2) (Butler et al. 2017) to perform multi-trait GBLUP. We modified the multi-trait GWAS model by removing the fixed marker effect and replacing the kinship matrix (𝐊) by the SNP-based GRM (𝐔_𝐒_) between DH lines.

### Cross-validation

To compare the prediction accuracy of different models and traits, we performed cross-validation. In all cross-validations, the training set comprised 250 DH lines and the remaining DH lines were assigned to the prediction set. Each cross-validation consisted of 100 random iterations. Prediction accuracy was calculated as the correlation between predicted genetic values and observed phenotypic values in the prediction set, divided by the square root of the trait heritability.

For each multi-trait model we compared two cross-validation scenarios: (1) MT-CV1, multi-trait models in which GDY from individual trials (target trait) and adjusted means across line *per se* trials of leaf rolling and leaf senescence (secondary traits) were included in the training set; and (2) MT-CV2, where phenotypes of the secondary traits were additionally provided in the prediction set.

## Results

### Sequencing-based genotyping

Shallow-seq of the 388 MAGIC DH lines resulted in an average depth of 5.7X. A total of 7.6 M SNPs with a missing rate less than 50% were retained from the 8.3 M SNPs identified in the eight founder lines. After filtering and imputation, a final SNP dataset containing 2,717,240 SNPs and no missing data was obtained for downstream analyses. Using the 600k array data as reference, the average genotyping error was estimated at 0.08%, comparable to the 0.04% observed in deep sequencing (50X) of the founder lines.

We assessed the genotyping efficiency of the shallow-seq by down-sampling the reads to six different depths (0.1 to 8X) using 31 DH lines (Fig. 1). Genotyping rate and genome coverage increased logarithmically with sequencing depth. Variance across samples was negligible. After filtering SNPs with 50% missing rate, SNP count rose sharply from 1X to 2X depth, with 6.2 M of the total 8.3 M SNPs retained at 2X (Fig. 1A). The genotyping error rate at positions supported by reads (unimputed dataset) remained consistent across depths. In contrast, the error rate of the imputed SNP dataset decreased drastically as depth increased, with 2X or higher achieving the same error rate as the raw dataset in which all SNP loci were supported by reads (Fig. 1B).

**Fig. 1.**
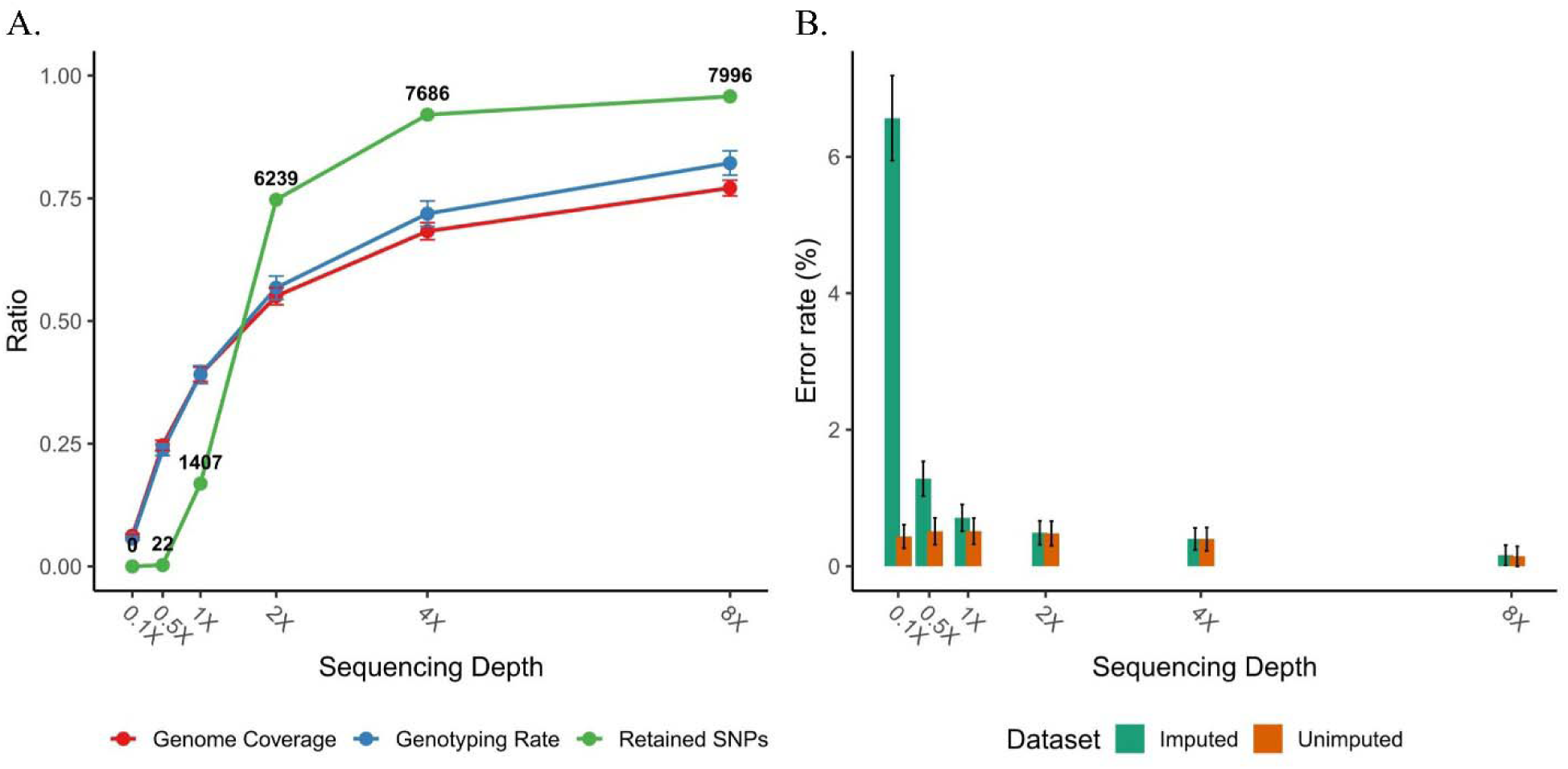
Efficiency and accuracy of shallow-seq. In both panels, the x-axis represents sequencing depth (0.1X to 8X). (A) Genotyping efficiency: Mean genome coverage (red), mean genotyping rate (blue), and the proportion of SNPs with < 0.5 missingness relative to the full set of 8.3 M SNPs (green). Numbers above the green points indicate the count of retained SNPs in thousands (k). (B) Genotyping error rate (%) for unimputed (brown) and imputed (green) SNP datasets, with standard deviation. Error bars represent standard deviations in both panels.

### Genomic characterization of the MAGIC population

The balanced intercrossing design minimized population structure in the MAGIC population. The first principal coordinate (PCo1) explained only 2.3% of the variance, with no obvious clustering among the three subfamilies. A neighbor joining (NJ) phylogenetic tree showed that the DH lines were distributed at equal distances from the root. LD decay ranged from 1.2 Mb to 3.3 Mb (*r^2^* = 0.2), with chromosome 10 exhibiting significantly longer LD compared to other chromosomes (Fig. S4).

Using the 2.7 million shallow-seq-based SNP markers, we reconstructed the mosaic of founder genomes in each MAGIC DH line. Fig. 2 illustrates the relative contribution of founder alleles along the ten chromosomes. The contributions of the eight founders are close to the expected value of 12.5%, consistent with balanced crossing. Along most chromosomal regions, founder contributions varied mildly. However, an unexpected pattern was observed on chromosome 9. Upon closer inspection, this pattern most likely resulted from the residual heterozygosity of inbred founder Lo1056. Thus, the plant sequenced to represent this founder genotype was polymorphic in a small region of chromosome 9 compared to those used for population development. Since this pattern was confined to a specific region, the material was retained for analysis.

**Fig. 2.**
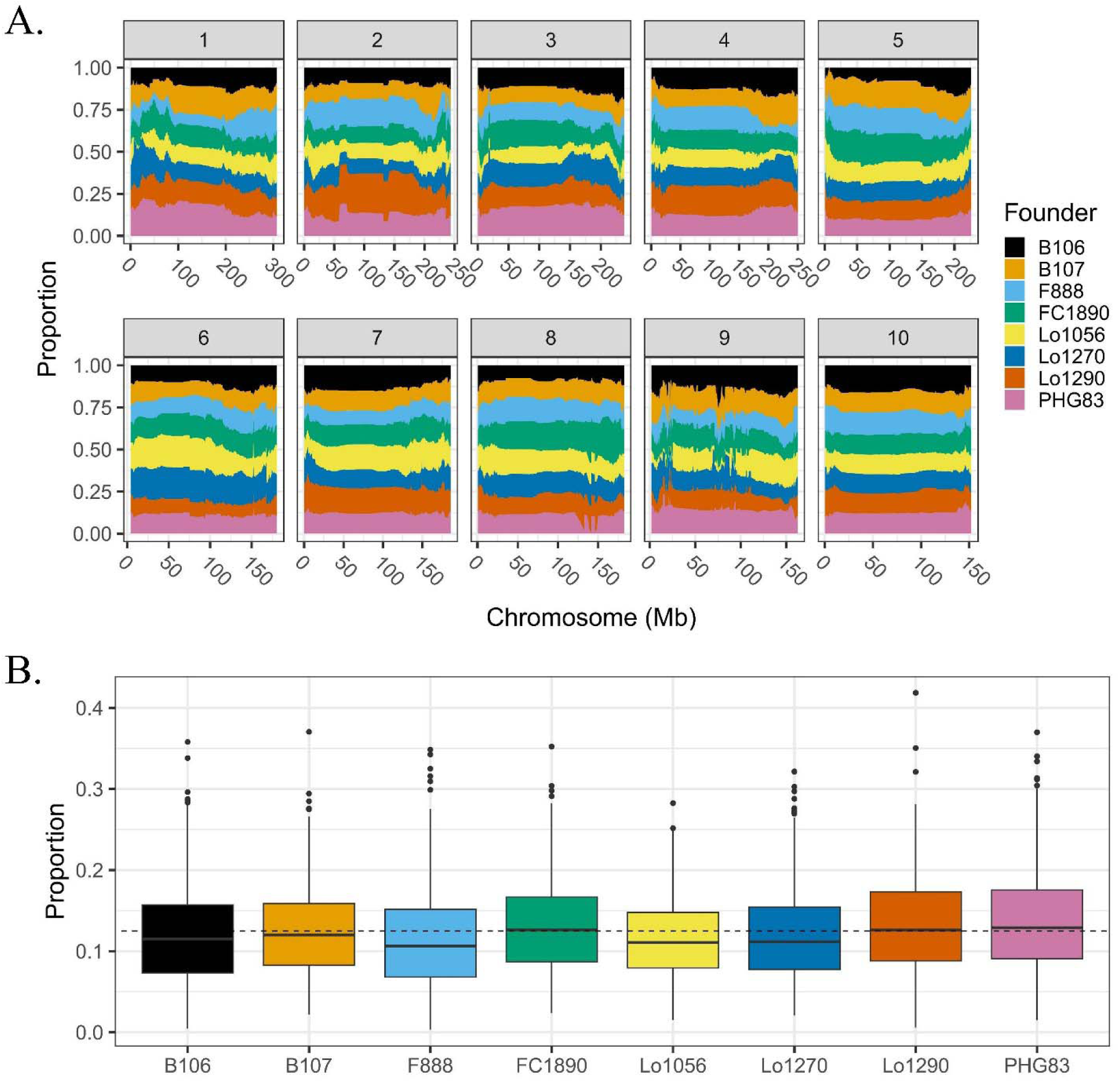
(A) Contributions of the eight founder lines to the MAGIC population along the 10 chromosomes. (B) Proportion of founder genome in the 388 DH lines.

On average, 3.8 recombination events were detected per chromosome, equivalent to 1.3 recombination events per meiosis (Fig. S5A). The estimation represents a lower bound as some recombination events occurring within IBD regions shared between founders or double crossovers remain undetected. The proportion of the genome in IBD between founder pairs ranged from 1% to 64%, with the maximum observed between Lo1056 and Lo1270, congruent with pedigree information that Lo1056 is the parent of Lo1270 (Fig. S5B).

Due to the IBD regions between founders, the assignment of founder alleles remained ambiguous in some genomic regions. On average, 31% of the genome across the 388 DH lines could not be assigned to any founder when applying a threshold of 0.9 on founder probability. By merging founder alleles into ancestral alleles, we observed an average of 5.9 ancestral alleles per locus as compared to eight founder alleles. Summing up the probabilities of the corresponding founder alleles to assign each ancestral allele reduced the unassigned proportion of the genome to 0.8%.

### Phenotypic characterization of the MAGIC population

Grain yield was measured in seven testcross trials and four line *per se* trials with average yield ranging from 4.8 to 10.1 t/ha and 1.3 to 3.8 t/ha, respectively (Fig. 3). In 2020, high rainfall at the test sites resulted in high yield in both irrigated and rainfed trials, representing optimal conditions. In 2021, high temperature and severe water stress occurred in Hungary clearly differentiating the two water regimes in Murony (MUR). The two German rainfed test sites (DTH, PNF) showed low to intermediate performance for grain yield. In 2022, line *per se* trials in Hungary and at one German site experienced stress conditions (MUR, DTH) leading to highly skewed yield distributions while growing conditions at the other two German sites (ROG, MER) were optimal for maize.

**Fig. 3.**
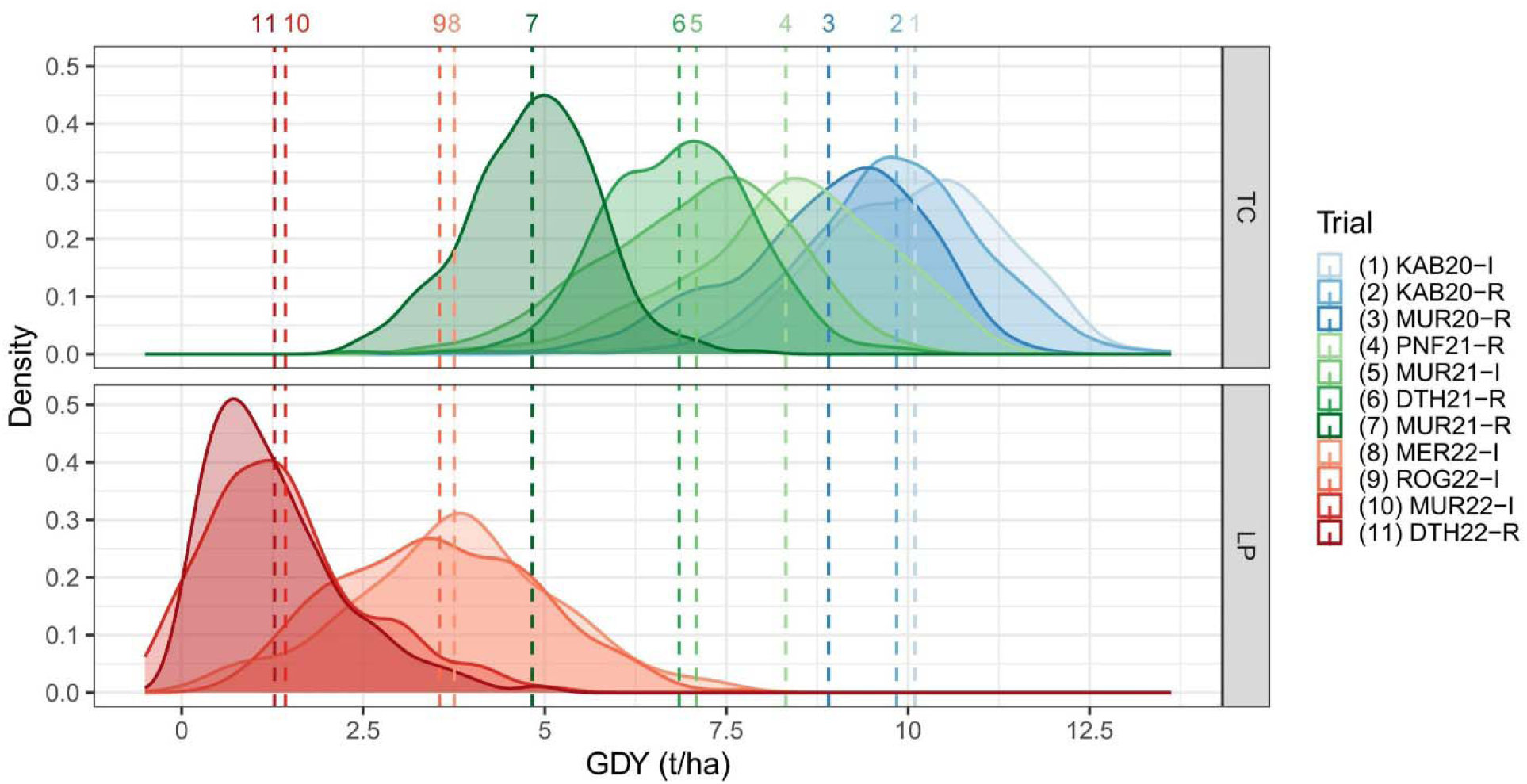
Density distribution of grain dry matter yield (GDY) based on adjusted entry means of 388 MAGIC DH lines evaluated as testcrosses (TC, upper panel) and lines *per se* (LP, lower panel) in seven (TC) and four (LP) trials. The vertical dashed lines indicate population means. Colors denote years: blue (2020), green (2021), and red (2022). Trial legends denote location, year, management (I = irrigated; R = rainfed), and yield rank in brackets.

As summarized in Table S3, pronounced genotype-by-trial interactions were observed for grain yield, with the G×T variance component ( 𝜎^2^𝑔𝑡) slightly exceeding the genetic component ( 𝜎^2^𝑔) with 𝜎^2^𝑔𝑡 : 𝜎^2^𝑔 ratios of 1.03 and 1.14 in testcross and line *per se* trials, respectively. Among the three proxy traits in line *per se* trials, leaf senescence showed the highest G×T ratio (1.02), followed by leaf rolling (0.78) and anthesis-silking interval (0.42). For testcross trials, the ratios were 0.88 for leaf senescence (based on two trials) and 0.72 for anthesis-silking interval, while leaf rolling was only scored in a single trial. Agronomic traits like final plant height and flowering time showed ratios below 0.3 for both testcross and line *per se* trials.

Pairwise phenotypic correlations among trials are shown in Fig. S6. Correlations for grain yield ranged from 0.06 to 0.79 among testcross trials and from 0.06 to 0.54 among line *per se* trials. As expected, correlations between testcross and line *per se* trials were generally low and not significant for grain yield. The three proxy traits displayed moderate correlations between trials, and significant correlations were observed between testcrosses and line *per se* performance. The agronomic traits final plant height and female flowering time showed high and consistent correlations across trials and between testcrosses and lines *per se*.

### Trait correlations between grain yield and the three proxy traits

The trials in Murony in 2021, where high temperatures and water stress occurred (Fig. S2), allow a closer look at correlations between grain yield and proxy traits with rainfed and irrigated trials side by side. Testcross grain yield decreased by 32% in the rainfed trial compared to the irrigated trial. In the rainfed trial, grain yield showed significant correlations with both leaf senescence (−0.25) and leaf rolling (−0.27), suggesting that delayed senescence and reduced leaf rolling were associated with higher yield (Fig. S7A). In particular, the top 10% genotypes exhibited greater stay-green (delayed senescence) and minimal leaf rolling than the remaining lines (Fig. S7B). In contrast, the anthesis-silking interval showed no significant correlation with grain yield.

We also assessed the correlations of the proxy traits evaluated on a line *per se* basis with testcross grain yield (Table 1). Leaf senescence measured across line *per se* rainfed trials showed negative correlations with testcross grain yield, reaching significance in trials at Murony (MUR), where temperatures were elevated. Leaf rolling followed a similar pattern but with weaker correlations. In contrast, anthesis-silking interval showed no consistent trend across testcross trials.

**Table 1.**
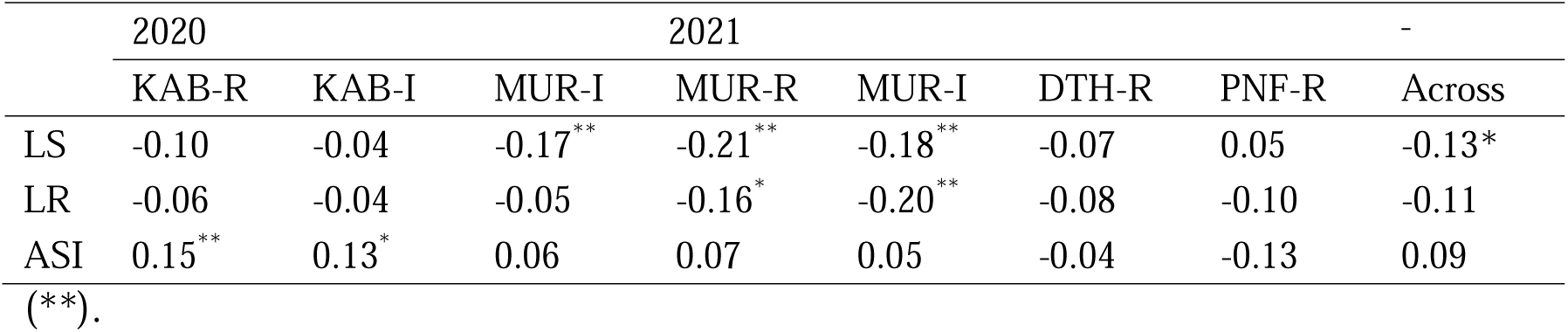
Pearson’s correlation coefficients between grain yield performance from seven testcross trials (and across trials) and the three proxy traits, leaf senescence (LS), leaf rolling (LR) and anthesis-silking interval (ASI), averaged across all line *per se* rainfed trials. Asterisks indicate significance levels after multiple testing adjustment: *p* < 0.05 (*), *p* < 0.01

### Overview of QTL detection

GWA analyses were conducted on adjusted entry means from individual trials and across trials. Table 2 summarizes the QTL sets obtained for grain yield and the three proxy traits in multi-trial analyses (see Table S4 for all traits). For testcross yield, 22 QTL were retained out of 71 single-trial associations, together explaining 45% of the genetic variance. Founder (FH) and ancestral (AH) haplotype-based mapping detected ten and seven QTL, respectively, with six and five overlapping with QTL regions detected by SNPs (see File S2 for the complete QTL list). For line *per se* grain yield, no QTL was detected using SNPs, while FH and AH methods identified 1 and 4 QTL, with no overlap between them.

**Table 2.** Summary of QTL for grain dry mater yield (GDY) and the three proxy traits, leaf senescence (LS), leaf rolling (LR), anthesis-silking interval (ASI). **nTrial** refers to the number of trials. Columns **SNP**, **FH**, and **AH** indicate the number of QTL identified by each method. Numbers in parentheses represent total number of QTL detected in individual trials. The proportion of genetic variance explained by all QTL identified using the SNP method is given by 𝝈^𝟐^ **explained**. **\***Testcross leaf rolling was measured in a single trial; therefore, only individual-

| | | nTrial | SNP | FH | AH | $\sigma_g^2$ explained |
| --- | --- | --- | --- | --- | --- | --- |
| Testcross | GDY | 7 | 22 (71) | 10 (24) | 7 (22) | 0.45 |
|  | LS | 2 | 13 (20) | 5 (6) | 5 (6) | 0.45 |
|  | *LR | 1 | - (5) | - (2) | - (2) | - (0.50) |
|  | ASI | 7 | 9 (13) | 2 (3) | 2 (4) | 0.53 |
| Line *per se* | GDY | 4 | 0 (0) | 1 (1) | 4 (6) | 0 |
|  | LS | 6 | 24 (36) | 5 (9) | 4 (7) | 0.53 |
|  | LR | 5 | 22 (47) | 7 (17) | 7 (16) | 0.30 |
|  | ASI | 13 | 15 (77) | 3 (26) | 1 (27) | 0.70 |

trial QTL counts are reported.

Fig. 4 presents the allelic effects of the 22 QTL discovered for testcross grain yield. Five QTL had consistent effects across all seven trials, while 17 QTL exhibited significant QxT interaction. The main effects at the consistent QTL ranged from 0.27 to 0.42 t/ha, with *qGDY(TC)08A* on chromosome 8 having the strongest effect. This QTL was also identified by both IBD-based methods, showing that PHG83 and Lo1290 carried the favorable alleles for yield improvement (Fig. S8A). For QTL with significant QxT interaction, response patterns varied. For instance, the QTL on chromosome 2 (*02A*) and 7 (*07A*) displayed opposite effects under optimal and stress conditions; the QTL on chromosome 1 (*01A*) had no effect under stress but under optimal conditions, whereas the QTL on chromosome 9 (*09B*) had a pronounced effect mainly in trials at Murony 2021, which experienced strong heat and drought stress.

**Fig. 4.**
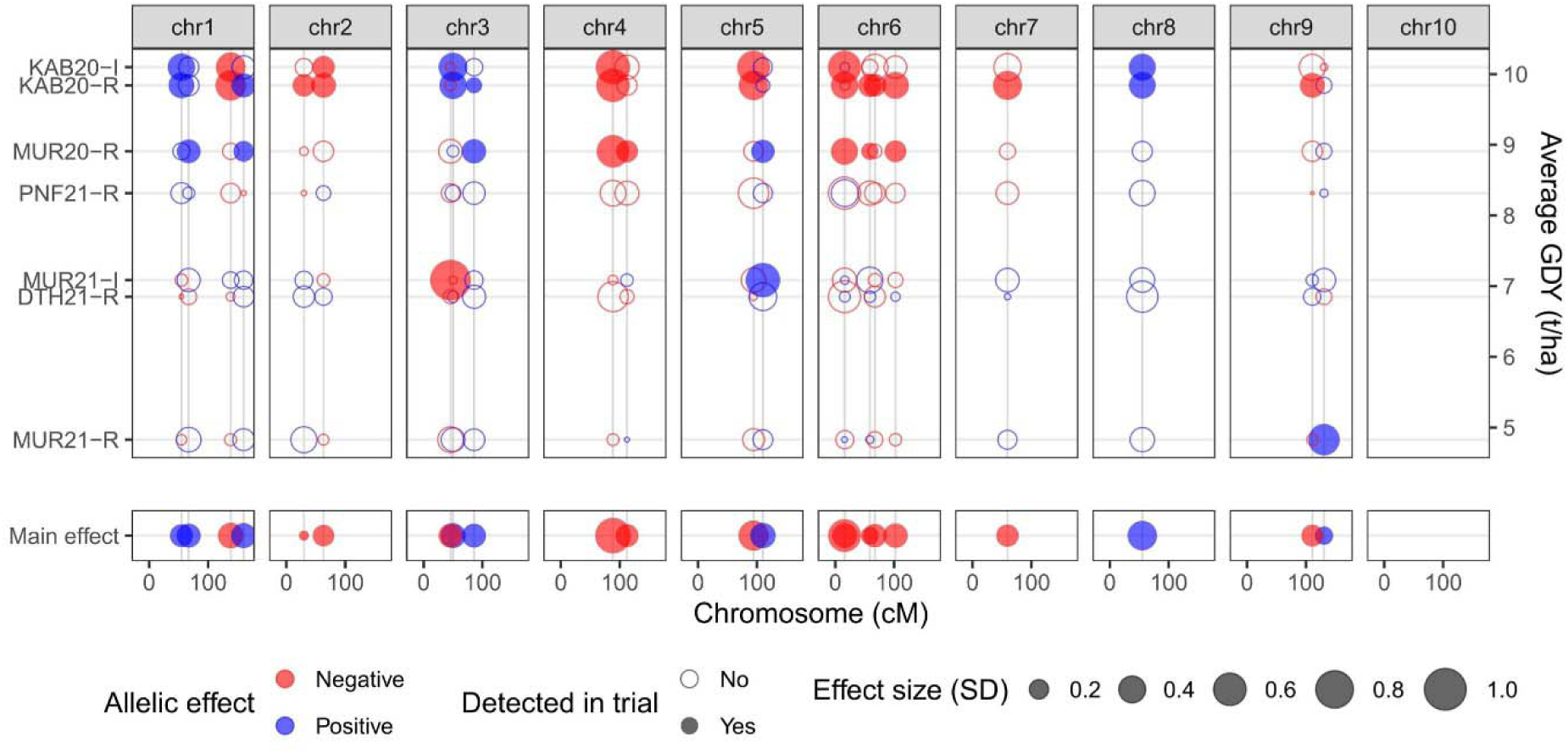
Grain yield QTL effects across individual testcross trials (upper panel) and combined main effect (lower panel). The x-axis shows the QTL position on the linkage map and the y-axis displays individual trials ordered by their average grain dry matter yield (GDY). Dot color indicates the direction of the allelic effect when the reference allele is substituted with the alternative allele. Solid or open circles differentiate significant from non-significant QTL in a given trial. Circle size reflects the magnitude of the QTL effect in that trial.

For proxy traits, 13 QTL were detected for leaf senescence and seven for anthesis-silking interval, explaining 45% and 53% of the genetic variance in testcross trials, respectively. Five QTL for leaf rolling were identified from a single testcross trial, accounting for 50% of the genetic variance (Table 2). In line *per se* trials, 24, 22, and 15 QTL were identified for the three traits, explaining 53%, 30%, and 70% of the genetic variance, respectively. Among the proxy trait QTL in line *per se* trials, four leaf senescence, two leaf rolling, and four anthesis-silking QTL overlapped with grain yield QTL intervals. However, the lead SNPs for proxy traits and grain yield were not in tight LD (maximum *r^2^* = 0.59), suggesting that they are unlikely to be the same QTL with pleiotropic effects.

We fitted a bivariate multi-trait model incorporating grain yield (across testcross trials) and one of the proxy traits (across line *per se* trials) to jointly estimate SNP effects for both traits. For the lead SNPs of QTL detected for leaf senescence and leaf rolling, effect estimates were significantly correlated with effects for grain yield at the same marker position (Fig. 5). This was not the case for the anthesis-silking interval QTL. A similar pattern was observed for the lead SNPs of the grain yield QTL (Fig. S9). Aligning with phenotypic correlations, alleles reducing leaf senescence and leaf rolling generally increased grain yield, and vice versa. Notably, QTL *qLS(LP)01C* (leaf senescence) and *qLR(LP)03C* (leaf rolling), displayed consistent effects across trials and had a substantial influence on testcross grain yield (Fig. 5), marking them as promising targets for further investigation.

**Fig. 5.**
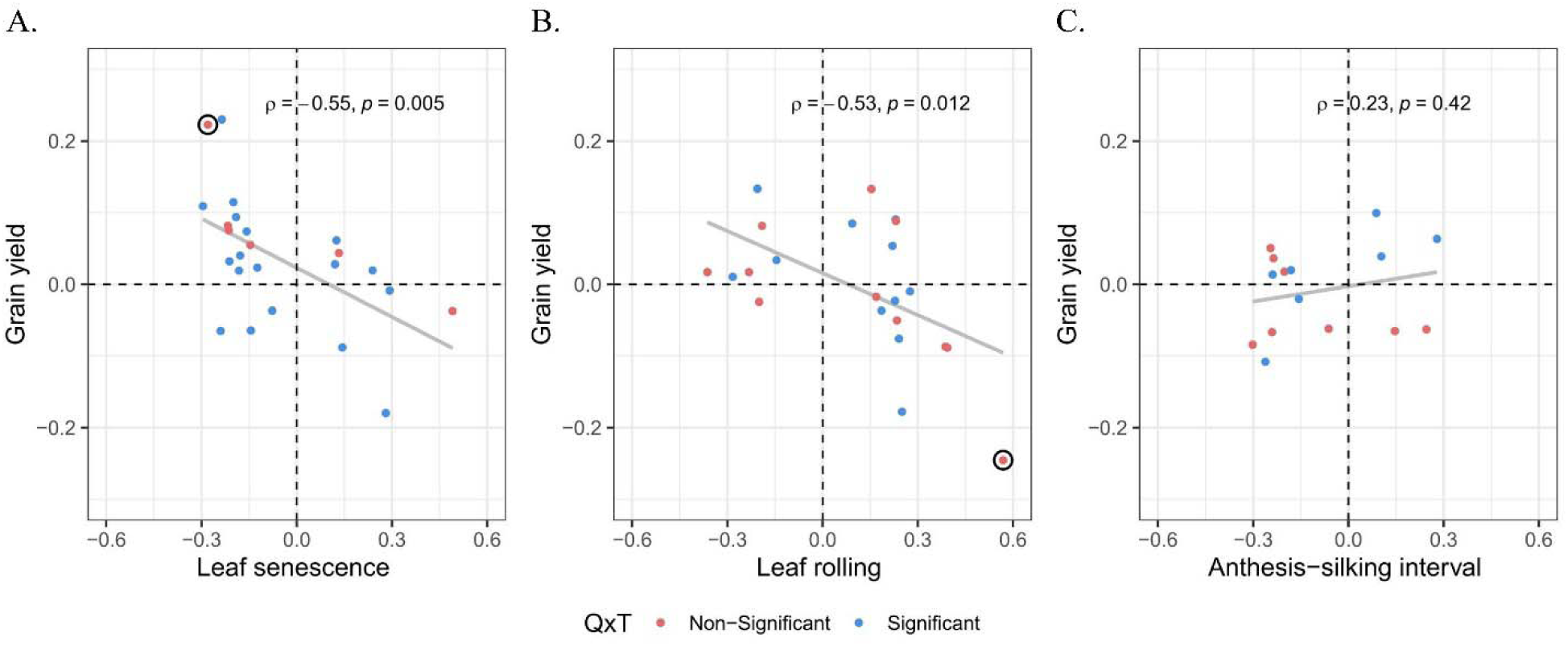
Correlation of SNP effects between grain yield and proxy traits at genomic positions of lead SNPs identified for proxy trait QTL. The y-axis shows SNP effect estimates on grain yield across all testcross trials, while the x-axis shows effects of the same SNP on (A) leaf senescence and (B) leaf rolling and (C) anthesis-silking interval across all line *per se* trials. Pearson correlation coefficients and associated significance levels are shown in each panel; Dot colors indicate whether the QTL exhibits significant QTL-by-Trial (QxT) interaction. Two environment-consistent QTL (non-significant QxT) with substantial effects on grain yield, *qLS(LP)01C* and *qLR(LP)03C*, are highlighted.

### Prediction accuracies for trait performance across trials

We evaluated accuracies of whole genome-based prediction for grain yield, the three proxy traits, and agronomic traits in the MAGIC population using SNP-based GBLUP (Table S5). Accuracies ranged from 0.33 (LS) to 0.74 (GDC) in testcross trials and 0.40 (LS) to 0.61 (FF) in line *per se* trials. Testcross yield could be predicted with intermediate accuracy (0.60). Among the three proxy traits measured in line *per se* trials, leaf rolling had the highest accuracy (0.60), followed by anthesis-silking interval (0.52) and leaf senescence (0.40).

We further investigated whether constructing genomic relationship matrices using IBD information (FH and AH) could enhance prediction accuracy. Elements of the matrices derived from FH and AH probabilities were significantly correlated (*r* = 0.91 and 0.94, respectively) with those of the SNP-based GRM. In terms of prediction accuracy, the AH method performed as well as the SNP method, with differences in accuracy ranging from - 0.02 to 0.01. The FH method was slightly inferior, with only 2 of 18 traits exceeding accuracies of the SNP method, and differences ranging from −0.04 to 0.01 (Table S5).

### Genomic prediction using leaf senescence and leaf rolling as secondary traits

We compared single-trait and multi-trait models to evaluate line *per se* leaf senescence and leaf rolling performance as secondary traits for genomic prediction of testcross grain yield. Using both traits as secondary traits in the MT-CV1 model (secondary traits available only for the training set) gave similar accuracies for prediction of testcross yield as the single-trait model. In MT-CV2 (secondary traits available in both training and prediction sets), slight improvements were observed in Murony 2021 trials, from 0.53 to 0.56 under rainfed and 0.47 to 0.51 under irrigated conditions. Accuracies in the other trials remained similar to the single-trait model. Analyzing the two proxy traits separately in MT-CV2 suggested that leaf senescence was the main driver of the observed improvement in Murony (Fig. 6).

**Fig. 6.**
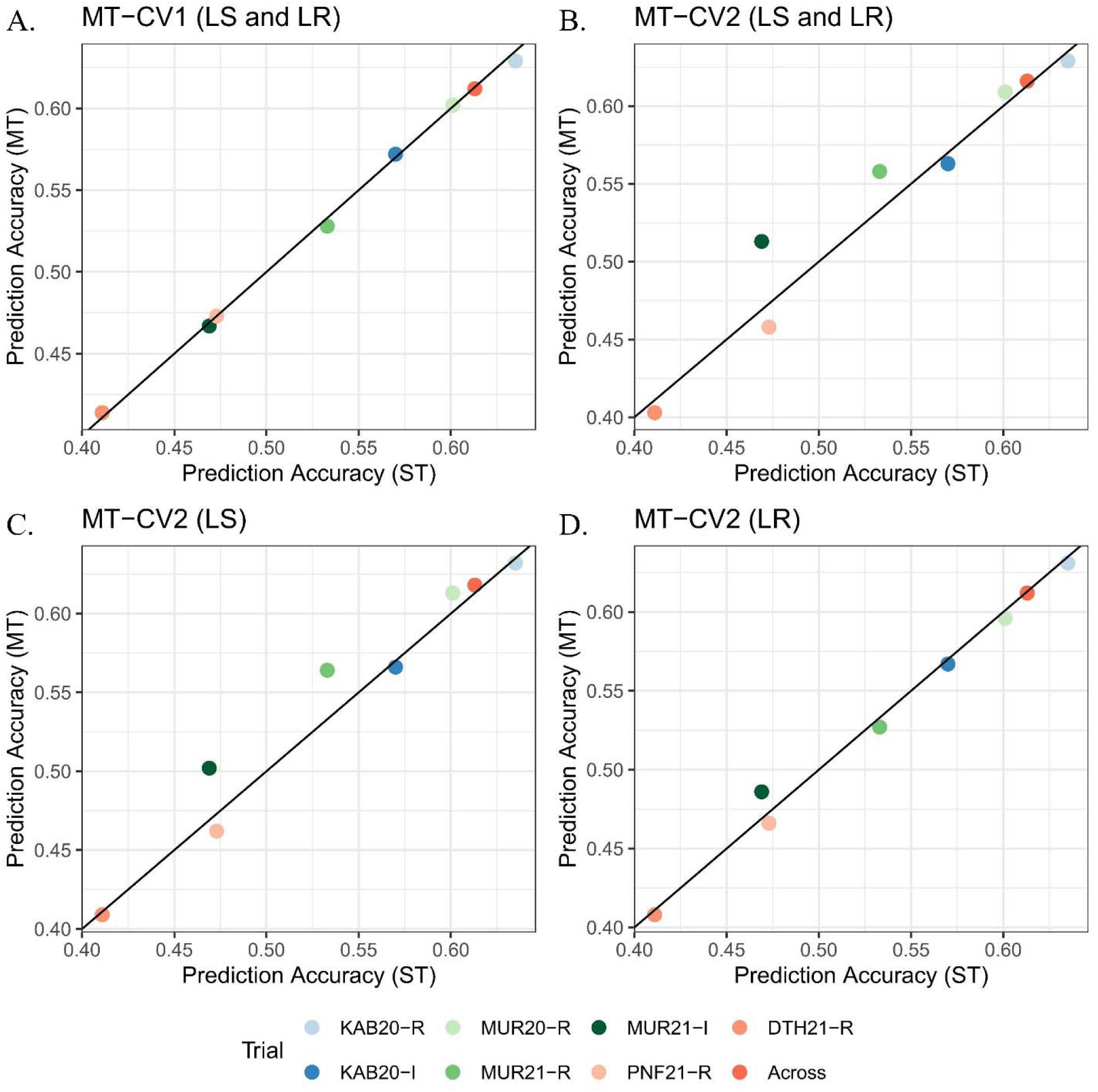
Comparison of grain yield prediction accuracies between single-trait (ST) and multi-trait (MT) models. MT models were evaluated when secondary traits were available only in the training set (CV1) or in both training and prediction sets (CV2). The x-axis shows ST prediction accuracies, and the y-axis shows MT prediction accuracies: (A) MT-CV1 using both leaf senescence (LS) and leaf rolling (LR) as secondary traits; (B) MT-CV2 using both LS and LR; (C) MT-CV2 using LS only; and (D) MT-CV2 using LR only. Colors denote seven individual trials and across trials. Values represent means of 100 cross-validation iterations.

## Discussion

Leveraging genetic diversity is essential for sustainable crop improvement. The DROPS panel (Millet et al. 2016), which has been characterized extensively across a wide range of European environments, represents a valuable resource of dent maize diversity. To further harness the genetic variation observed for our target traits, we selected eight inbred lines with contrasting yield performance under drought and heat stress, ensuring limited variation in flowering time, to develop a MAGIC population. This population was genotyped using whole-genome shallow sequencing and phenotyped across 21 field experiments. We dissected the genetic architecture of important agronomic traits, primarily grain yield, evaluated the usefulness of proxy traits for improving grain yield under stress conditions, and provided recommendations for optimizing MAGIC genotyping and mapping strategies.

### Evaluating proxy traits for yield improvement

The eight MAGIC founders were selected to represent variation in heat and drought tolerance and to capture part of the genetic diversity of the Non-Stiff Stalk heterotic group within the DROPS panel. They exhibited polymorphism for 74% of the markers analyzed in the DROPS panel and were polymorphic for 39 of the 48 grain yield QTL identified (Millet et al. 2016). In our seven testcross trials, the average yield range for the MAGIC population (4.8 to 10.1 t/ha) covered the broader environmental variation observed across the 29 experiments of the DROPS panel (1.5 to 11.2 t/ha) only partially. Despite capturing only a subset of the genetic diversity and a narrower environmental range, seven of the 48 DROPS grain yield QTL overlapped with the genomic positions of grain yield QTL detected in this study (Table S6).

Beyond testcross yield, we evaluated three long-established drought proxy traits, leaf senescence, leaf rolling and anthesis-silking interval in the MAGIC population (Bänziger et al. 2000). These traits have been incorporated into selection indices for improving yield in drought-prone environments in testcross trials (Edmeades et al. 1999). In addition to testcross evaluations, we also assessed these traits in line *per se* trials, which can be conducted in short observation plots and facilitate scalable assessment when combined with UAV-based high-throughput phenotyping (Baret et al. 2018; Cerrudo et al. 2017). This scalability may potentially compensate for the generally low correlation observed between the proxy traits and grain yield. Here, we observed modest but significant correlations between grain yield and both leaf senescence and leaf rolling. The low observed correlations align with Monneveux et al. (2008), who reported the marginal relevance of conventional secondary traits for direct selection. However, despite low overall correlations, leaf senescence and leaf rolling remain distinguishing features of tolerant lines, as the top 10% of yielding genotypes in our MAGIC population exhibited significantly delayed senescence and reduced leaf rolling. The negative correlation between leaf rolling and yield suggests that rolling reflects drought sensitivity rather than a protective mechanism in these genotypes (Parajuli et al. 2018). The lack of correlation of the anthesis-silking interval with grain yield in testcrosses was likely due to the short interval under stress condition (2.3 days in MUR21-R), indicating that this trait is already optimized in modern hybrid maize (Araus et al. 2012).

In line *per se* trials, leaf senescence and leaf rolling were correlated with testcross yield only under high-temperature and water-deficit conditions. Although no major pleiotropic QTL were identified, a clear trend emerged among the detected proxy trait QTL: alleles associated with delayed senescence and reduced rolling generally showed positive effects on testcross grain yield, indicating that accumulating these favorable alleles could enhance yield performance. A shared polygenic basis, pleiotropy of minor genes, is further supported by the improved prediction accuracy of grain yield when these two traits were included in the multi-trait GBLUP models. However, given the relatively small gains in prediction accuracy, the additional efforts required for including proxy traits in a breeding context seem justified only if they can be assessed in high throughput and with high precision.

### QTL for trait improvement

In this study, we report several QTL for grain yield, proxy traits and agronomic traits, which represent valuable targets for trait improvement. Sekhon et al. (2019) reported 64 candidate genes for leaf senescence, via a comprehensive GWAS and gene expression experiment using a diversity panel composed primarily of Stiff Stalk and Non-Stiff Stalk dent lines. We found four of these genes located within our QTL regions (File S2): *Zm00001d030968* (Flowering locus K domain protein), *Zm00001d052798* (Trihelix-transcription factor 3), *Zm00001d002967* (AFG1-like ATPase family protein), *Zm00001d020396* (Trehalose-6-phospate synthase13, *trps13*). Thus, our results provide a valuable resource for a future study on potential candidate genes in these genomic regions.

For leaf rolling, one QTL (*qLR(LP)03C*) is of particular interest due to its potential pleiotropic effects on yield and its consistent effect across trials. This QTL was detected by all three QTL detection methods. The Lo1056 and B106 alleles reduced leaf rolling (beneficial) with the Lo1056 allele showing the stronger effect (Fig. 8B). This leaf rolling QTL co-localized with a leaf angle QTL identified in the maize NAM population (Tian et al. 2011), suggesting a potential genetic link between the leaf rolling mechanism and overall leaf architecture.

While the anthesis-silking interval was not correlated with testcross grain yield, a significant correlation was observed with line *per se* grain yield (data not shown). The beneficial allele which shortens ASI in line *per se* trials therefore might remain important for seed production of inbred lines.

Among agronomic trait QTL, a plant height QTL, *qPH_final(LP)01A*, showed a strong effect (∼11 cm) across line *per se* trials but not in testcross trials. The B106 allele significantly reduced plant height. A candidate gene, *Brachytic2* (*Br2*) (Multani et al. 2003), involved in polar auxin transport, lies ∼2.2 Mb away from the lead SNP of the QTL. Sequencing data revealed a potential 1 kb duplication within exon 5 of *Br2* in the B106 allele, which may contribute to the reduction in plant height. As the *Br2* gene has been exploited to develop short-stature maize hybrids, marketed as “smart corn”, the B106 allele might also represent a valuable allele for future breeding applications.

### IBS and IBD-based methods for QTL detection

The known founder genotypes of the MAGIC population enabled genomic mosaic reconstruction and modeling of multi-allelic QTL effects (Li et al. 2021). Odell et al. (2022) recommended applying multiple approaches to obtain comprehensive and complementary results. SNPs have long been used in multi-trait and -environment QTL analysis (Jiang and Zeng 1995; Korte et al. 2012). More recently, this framework has been extended to multi-allelic (FH and AH) approaches. Garin et al. (2020) modeled QTL effects as fixed effects, decomposing them into main effects and Q×E interaction effects. Alternatively, Li et al. (2024) treated QTL effects as random, allowing variances to differ among environments and reduce complexity. Fixed models might lose power as the number of environments or trials increases. The assumption of normally distributed allelic effects in the mixed model might not always reflect biological reality. Here, we applied IBD-based QTL mapping (FH and AH) with a random effect model and compared it with the SNP-based approach. The SNP method identified a greater number of unique QTL in both individual trial and multi-trial analyses. On the other hand, individual QTL explained more phenotypic variance with IBD-based methods, likely due to the estimation of multiple allelic effects; however, the total QTL variance explained per trait was greater with the SNP-based approach (Fig. S10). Given the dense set of 2.7 million SNPs, which likely approaches saturation for capturing all chromosome recombination events, and the broader QTL set identified, our analyses focused primarily on SNP-based results, with IBD-based methods as a complement.

Based on these results, we recommend combining SNP-based and IBD-based (FH or AH) methods for MAGIC populations. IBD-based methods are ideal for downstream gene cloning, as they identify fewer but stronger QTL, are less sensitive to individual SNP genotyping errors, and provide founder allelic effects. Furthermore, at low marker densities, they show advantages: when SNPs were reduced to 600k and 15k array positions (Method S2), LOD curves from the IBD-based methods remained largely unchanged, whereas SNP detected fewer QTL with lower significance for lead SNPs in the 15k dataset (Fig. S11). In contrast, at sufficiently high SNP densities, SNP-based QTL detection offers greater power for detection of QTL with few segregating alleles, and is preferable for marker-assisted selection due to the well-established frameworks for marker design, effect estimation, and statistical modeling.

### Shallow-seq efficiency

SNP marker density and precision are determined by the genotyping platform employed. Reduced-representation sequencing (RRS), which targets only a small fraction of the genome for sequencing, and fixed-content SNP arrays have been widely used in maize studies due to their high accuracy, low missingness, comparability across experiments, and reasonable costs (Rasheed et al. 2017). These platforms are well suited for genomic prediction and linkage map construction (Chen et al. 2014; Crossa et al. 2013; Ganal et al. 2011). However, both approaches are limited to fixed genomic positions and prone to ascertainment bias, making them less effective for discovering rare variants in diverse germplasm panels (Davey et al. 2011; Romay et al. 2013). Low-coverage whole-genome sequencing (lcWGS), referred to here as shallow-seq, combined with effective imputation, offers more uniform genome-wide coverage and higher marker density (Scheben et al. 2017). With sequencing costs continuing to decline, lcWGS is a practical and scalable option for maize genetics (Lou et al. 2021). The drawbacks of this technology are higher error rates, greater missingness, and limited comparability across experiments, but these disadvantages can be mitigated by optimizing sequencing depth.

The optimal sequencing depth can differ depending on population type, founder information, minor allele frequency, and sample heterozygosity (Lou et al. 2021). The rich body of sequencing data generated for the MAGIC (>50X for the founders and average 5.7X for the DH lines) provided a valuable resource for evaluating and refining genotyping pipelines. Here, we evaluated a scenario with known founder genotypes and examined the number of SNPs detected across sequencing depths. Our results showed that 2X is a turning point, beyond which gains in SNP detection, genotyping rate, and genome coverage begin to plateau. At this depth, 75% of known SNP loci were retained with a missing rate below 0.5, allowing for effective imputation. Given the fixed costs of library preparation, 2X offered the best balance of cost and performance. However, when budgets are limited, maintaining at least 0.5X coverage per sample remains a practical alternative, still yielding over 22K SNPs, sufficient for IBD-based mapping in this population (Fig. S11).

Progressing from the dent diversity panel for crop improvement, we developed a MAGIC population supported by extensive genotypic and phenotypic datasets. This study focused primarily on dissecting the genetic architecture of grain yield and evaluating the utility of drought-related proxy traits for yield improvement. Beyond this, the MAGIC resource allowed to address a broad range of research questions such as optimizing lcWGS and founder detection methods, serving as a training population for marker-based genomic selection, and candidate gene identification for agronomic traits.

## Author Contribution Statement

CCS, MM, MO and TPr conceived and designed the study. Plant material preparation and development was performed by AH, MO, CU and TPr, data collection was performed by YCL, SU, AH. Genotypic data were produced by TP and CU and processed by YCL. YCL and AS analyzed the phenotypic and genotypic data under the supervision of CCS. TPo developed software for genotypic data analysis. CCS and MO acquired funding for the study. The first draft of the manuscript was written by YCL. CCS edited the manuscript. All authors commented on previous versions of the manuscript. All authors read and approved the final manuscript.

## Acknowledgement

We thank the Plant Technology Center (Technical University of Munich, Germany) for providing infrastructure and technical support during field experiments. We are grateful to the technical staff of KWS SAAT SE & Co. KGaA, Technical University of Munich and the student employees of the chair of plant breeding at Technical University of Munich for their support and thorough phenotypic data collection required for this study. The study was funded by the Federal Ministry of Education and Research (BMBF, Germany) within the scope of the funding initiative “Plant Breeding Research of the Bioeconomy” (Funding ID: 031B0195 and 031B0882; www.europeanmaize.net).

## Compliance with ethical standards

### Ethical standards

The authors declare that this study complies with the current laws of the countries in which the experiments were performed.

## Availability of data and materials

All data and materials are available through material transfer agreements upon request.

Raw WGS reads have been deposited in NCBI BioProjects PRJNA1434727. The curated phenotypic data (including raw plot-level observations and adjusted entry means) and SNP variant calling files (VCFs) are available via Figshare at the following URL: https://doi.org/10.6084/m9.figshare.31557775.

## Supplementary material

MAGICpaper_TAG_Supplementary_MethodFigureTable.pdf MAGICpaper_TAG_FileS1_WeatherData.xlsx MAGICpaper_TAG_FileS2_QTLResults.xlsx

## References

Araus JL, Serret MD, Edmeades GO (2012) Phenotyping maize for adaptation to drought. Front Physiol 3:305 10.3389/fphys.2012.00305

Astle W, Balding DJ (2009) Population Structure and Cryptic Relatedness in Genetic Association Studies. Stat Sci 24:451–471 10.1214/09-Sts307

Bänziger M, Edmeades GO, Beck D, Bellon MR (2000) Breeding for drought and nitrogen stress tolerance in maize: from theory to practice. CIMMYT, Mexico

Baret F, Madec S, Irfan K, Lopez J, Comar A, Hemmerlé M, Dutartre D, Praud S, Tixier MH (2018) Leaf-rolling in maize crops: from leaf scoring to canopy-level measurements for phenotyping. J Exp Bot 69:2705–2716 10.1093/jxb/ery071

Benjamini Y, Hochberg Y (1995) Controlling the False Discovery Rate: A Practical and Powerful Approach to Multiple Testing. J R Stat Soc B 57:289–300 10.1111/j.2517-6161.1995.tb02031.x

Bolger AM, Lohse M, Usadel B (2014) Trimmomatic: a flexible trimmer for Illumina sequence data. Bioinformatics 30:2114–2120 10.1093/bioinformatics/btu170

Bonfield JK, Marshall J, Danecek P, Li H, Ohan V, Whitwham A, Keane T, Davies RM (2021) HTSlib: C library for reading/writing high-throughput sequencing data. Gigascience 10 10.1093/gigascience/giab007

Browning BL, Zhou Y, Browning SR (2018) A One-Penny Imputed Genome from Next-Generation Reference Panels. Am J Hum Genet 103:338–348 10.1016/j.ajhg.2018.07.015

Bruce WB, Edmeades GO, Barker TC (2002) Molecular and physiological approaches to maize improvement for drought tolerance. J Exp Bot 53:13–25 10.1093/jexbot/53.366.13

Butler D, Cullis B, Gilmour A, Gogel B, Thompson R (2017) ASReml-R reference manual version 4. VSN International Ltd, Hemel Hempstead, HP1 1ES, UK

Caicedo M, Munaiz ED, Malvar RA, Jiménez JC, Ordas B (2021) Precision Mapping of a Maize MAGIC Population Identified a Candidate Gene for the Senescence-Associated Physiological Traits. Front Genet 12:716821 10.3389/fgene.2021.716821

Cerrudo D, González Pérez L, Mendoza Lugo JA, Trachsel S (2017) Stay-Green and Associated Vegetative Indices to Breed Maize Adapted to Heat and Combined Heat-Drought Stresses. Remote Sens-Basel 9:235 10.3390/rs9030235

Chen ZL, Wang BB, Dong XM, Liu H, Ren LH, Chen J, Hauck A, Song WB, Lai JS (2014) An ultra-high density bin-map for rapid QTL mapping for tassel and ear architecture in a large F_2_ maize population. Bmc Genomics 15:433 10.1186/1471-2164-15-433

Crossa J, Beyene Y, Kassa S, Pérez P, Hickey JM, Chen C, de los Campos G, Burgueño J, Windhausen VS, Buckler E, Jannink JL, Cruz MAL, Babu R (2013) Genomic Prediction in Maize Breeding Populations with Genotyping-by-Sequencing. G3-Genes Genom Genet 3:1903–1926 10.1534/g3.113.008227

Danecek P, Auton A, Abecasis G, Albers CA, Banks E, DePristo MA, Handsaker RE, Lunter G, Marth GT, Sherry ST, McVean G, Durbin R, Grp GPA (2011) The variant call format and VCFtools. Bioinformatics 27:2156–2158 10.1093/bioinformatics/btr330

Davey JW, Hohenlohe PA, Etter PD, Boone JQ, Catchen JM, Blaxter ML (2011) Genome-wide genetic marker discovery and genotyping using next-generation sequencing. Nat Rev Genet 12:499–510 10.1038/nrg3012

Dell’Acqua M, Gatti DM, Pea G, Cattonaro F, Coppens F, Magris G, Hlaing AL, Aung HH, Nelissen H, Baute J, Frascaroli E, Churchill GA, Inzé D, Morgante M, Pè ME (2015) Genetic properties of the MAGIC maize population: a new platform for high definition QTL mapping in *Zea mays*. Genome Biol 16:167 10.1186/s13059-015-0716-z

Edmeades GO, Bolaños J, Chapman SC, Lafitte HR, Bänziger M (1999) Selection improves drought tolerance in tropical maize populations: I. Gains in biomass, grain yield, and harvest index. Crop Science 39:1306–1315 10.2135/cropsci1999.3951306x

Effendi R, Priyanto SB, Aqil M, Azrai M (2019) Drought adaptation level of maize genotypes based on leaf rolling, temperature, relative moisture content, and grain yield parameters. Iop C Ser Earth Env 270:012016 10.1088/1755-1315/270/1/012016

Ferguson JN, Caproni L, Walter J, Shaw K, Arce-Cubas L, Baines A, Thein MS, Mager S, Taylor G, Cackett L, Mathan J, Vath RL, Martin L, Genty B, Pè ME, Lawson T, Dell’Acqua M, Kromdijk J (2025) A deficient CP24 allele defines variation for dynamic nonphotochemical quenching and photosystem II efficiency in maize. Plant Cell 37 10.1093/plcell/koaf063

Freyman WA, McManus KF, Shringarpure SS, Jewett EM, Bryc K, Auton A, Team MR (2021) Fast and Robust Identity-by-Descent Inference with the Templated Positional Burrows-Wheeler Transform. Mol Biol Evol 38:2131–2151 10.1093/molbev/msaa328

Ganal MW, Durstewitz G, Polley A, Bérard A, Buckler ES, Charcosset A, Clarke JD, Graner EM, Hansen M, Joets J, Le Paslier MC, McMullen MD, Montalent P, Rose M, Schön CC, Sun Q, Walter H, Martin OC, Falque M (2011) A Large Maize (Zea mays L.) SNP Genotyping Array: Development and Germplasm Genotyping, and Genetic Mapping to Compare with the B73 Reference Genome. Plos One 6 10.1371/journal.pone.0028334

Garin V, Malosetti M, van Eeuwijk F (2020) Multi-parent multi-environment QTL analysis: an illustration with the EU-NAM Flint population. Theor Appl Genet 133:2627–2638 10.1007/s00122-020-03621-0

Garrison E, Marth G (2012) Haplotype-based variant detection from short-read sequencing. arXiv preprint arXiv:12073907 10.48550/arXiv.1207.3907

Hallauer AR, Carena MJ, Miranda Filho Jd (2010) Quantitative genetics in maize breeding. Springer Science & Business Media

Hill WG, Weir BS (1988) Variances and Covariances of Squared Linkage Disequilibria in Finite Populations. Theor Popul Biol 33:54–78 10.1016/0040-5809(88)90004-4

Huang BE, Verbyla KL, Verbyla AP, Raghavan C, Singh VK, Gaur P, Leung H, Varshney RK, Cavanagh CR (2015) MAGIC populations in crops: current status and future prospects. Theor Appl Genet 128:999–1017 10.1007/s00122-015-2506-0

Hufford MB, Seetharam AS, Woodhouse MR, Chougule KM, Ou SJ, Liu JN, Ricci WA, Guo TT, Olson A, Qiu YJ, Della Coletta R, Tittes S, Hudson AI, Marand AP, Wei SR, Lu ZY, Wang B, Tello-Ruiz MK, Piri RD, Wang N, Kim DW, Zeng YB, O’Connor CH, Li XR, Gilbert AM, Baggs E, Krasileva KV, Portwood JL, Cannon EKS, Andorf CM, Manchanda N, Snodgrass SJ, Hufnagel DE, Jiang QH, Pedersen S, Syring ML, Kudrna DA, Llaca V, Fengler K, Schmitz RJ, Ross-Ibarra J, Yu JM, Gent JI, Hirsch CN, Ware D, Dawe RK (2021) De novo assembly, annotation, and comparative analysis of 26 diverse maize genomes. Science 373:655–+ 10.1126/science.abg5289

Jiang CJ, Zeng ZB (1995) Multiple-Trait Analysis of Genetic-Mapping for Quantitative Trait Loci. Genetics 140:1111–1127 10.1093/genetics/140.3.1111

Knapp SJ, Stroup WW, Ross WM (1985) Exact Confidence-Intervals for Heritability on a Progeny Mean Basis. Crop Science 25:192–194 10.2135/cropsci1985.0011183X002500010046x

Korte A, Vilhjálmsson BJ, Segura V, Platt A, Long Q, Nordborg M (2012) A mixed-model approach for genome-wide association studies of correlated traits in structured populations. Nat Genet 44:1066–+ 10.1038/ng.2376

Lee EA, Tollenaar M (2007) Physiological basis of successful breeding strategies for maize grain yield. Crop Science 47:S202–S215 10.2135/cropsci2007.04.0010IPBS

Li H, Durbin R (2009) Fast and accurate short read alignment with Burrows-Wheeler transform. Bioinformatics 25:1754–1760 10.1093/bioinformatics/btp324

Li H, Handsaker B, Wysoker A, Fennell T, Ruan J, Homer N, Marth G, Abecasis G, Durbin R, Proc GPD (2009) The Sequence Alignment/Map format and SAMtools. Bioinformatics 25:2078–2079 10.1093/bioinformatics/btp352

Li WH, Boer MP, Joosen RVL, Zheng CZ, Percival-Alwyn L, Cockram J, Van Eeuwijk FA (2024) Modeling QTL-by-environment interactions for multi-parent populations. Front Plant Sci 15 10.3389/fpls.2024.1410851

Li WH, Boer MP, van Rossum BJ, Zheng CZ, Joosen RVL, van Eeuwijk FA (2022) statgenMPP: an R package implementing an IBD-based mixed model approach for QTL mapping in a wide range of multi-parent populations. Bioinformatics 38:5134–5136 10.1093/bioinformatics/btac662

Li WH, Boer MP, Zheng CZ, Joosen RVL, van Eeuwijk FA (2021) An IBD-based mixed model approach for QTL mapping in multiparental populations. Theor Appl Genet 134:3643–3660 10.1007/s00122-021-03919-7

Lou RN, Jacobs A, Wilder AP, Therkildsen NO (2021) A beginner’s guide to low-coverage whole genome sequencing for population genomics. Molecular Ecology 30:5966–5993 10.1111/mec.16077

Mantel N (1967) The detection of disease clustering and a generalized regression approach. Cancer research 27:209–220

McCouch SR (1997) Report on QTL nomenclature. Rice Genet Newsl 14:11–13

Michel KJ, Lima DC, Hundley H, Singan V, Yoshinaga Y, Daum C, Barry K, Broman KW, Buell CR, de Leon N, Kaeppler SM (2022) Genetic mapping and prediction of flowering time and plant height in a maize Stiff Stalk MAGIC population. Genetics 221 10.1093/genetics/iyac063

Millet EJ, Kruijer W, Coupel-Ledru A, Prado SA, Cabrera-Bosquet L, Lacube S, Charcosset A, Welcker C, van Eeuwijk F, Tardieu F (2019) Genomic prediction of maize yield across European environmental conditions. Nat Genet 51:952–+ 10.1038/s41588-019-0414-y

Millet EJ, Welcker C, Kruijer W, Negro S, Coupel-Ledru A, Nicolas SD, Laborde J, Bauland C, Praud S, Ranc N, Presterl T, Tuberosa R, Bedo Z, Draye X, Usadel B, Charcosset A, Van Eeuwijk F, Tardieu F (2016) Genome-Wide Analysis of Yield in Europe: Allelic Effects Vary with Drought and Heat Scenarios. Plant Physiol 172:749–764 10.1104/pp.16.00621

Monneveux P, Sanchez C, Tiessen A (2008) Future progress in drought tolerance in maize needs new secondary traits and cross combinations. J Agr Sci-Cambridge 146:287–300 10.1017/S0021859608007818

Morgulis A, Gertz EM, Schäffer AA, Agarwala R (2006) A fast and symmetric DUST implementation to mask low-complexity DNA sequences. J Comput Biol 13:1028–1040 10.1089/cmb.2006.13.1028

Multani DS, Briggs SP, Chamberlin MA, Blakeslee JJ, Murphy AS, Johal GS (2003) Loss of an MDR transporter in compact stalks of maize *br2* and sorghum *dw3* mutants. Science 302:81–84 10.1126/science.1086072

Odell SG, Hudson A, Praud S, Dubreuil P, Tixier MH, Ross-Ibarra J, Runcie DE (2022) Modeling allelic diversity of multiparent mapping populations affects detection of quantitative trait loci. G3-Genes Genom Genet 12 10.1093/g3journal/jkac011

Paradis E, Schliep K (2019) ape 5.0: an environment for modern phylogenetics and evolutionary analyses in R. Bioinformatics 35:526–528 10.1093/bioinformatics/bty633

Parajuli S, Ojha B, Ferrara G (2018) Quantification of secondary traits for drought and low nitrogen stress tolerance in inbreds and hybrids of maize (Zea mays L.). J Plant Genet Breed, p 106

Pook T, Nemri A, Segovia EGG, Torres DV, Simianer H, Schoen CC (2021) Increasing calling accuracy, coverage, and read-depth in sequence data by the use of haplotype blocks. Plos Genet 17 10.1371/journal.pgen.1009944

Pook T, Schlather M, de los Campos G, Mayer M, Schoen CC, Simianer H (2019) HaploBlocker: Creation of Subgroup-Specific Haplotype Blocks and Libraries. Genetics 212:1045–1061 10.1534/genetics.119.302283

Purcell S, Neale B, Todd-Brown K, Thomas L, Ferreira MAR, Bender D, Maller J, Sklar P, de Bakker PIW, Daly MJ, Sham PC (2007) PLINK: A tool set for whole-genome association and population-based linkage analyses. Am J Hum Genet 81:559–575 10.1086/519795

Rasheed A, Hao YF, Xia XC, Khan A, Xu YB, Varshney RK, He ZH (2017) Crop Breeding Chips and Genotyping Platforms: Progress, Challenges, and Perspectives. Mol Plant 10:1047–1064 10.1016/j.molp.2017.06.008

Revell LJ (2012) phytools: an R package for phylogenetic comparative biology (and other things). Methods Ecol Evol 3:217–223 10.1111/j.2041-210X.2011.00169.x

Rida S, Maafi O, López-Malvar A, Revilla P, Riache M, Djemel A (2021) Genetics of Germination and Seedling Traits under Drought Stress in a MAGIC Population of Maize. Plants-Basel 10 10.3390/plants10091786

Romay MC, Millard MJ, Glaubitz JC, Peiffer JA, Swarts KL, Casstevens TM, Elshire RJ, Acharya CB, Mitchell SE, Flint-Garcia SA, McMullen MD, Holland JB, Buckler ES, Gardner CA (2013) Comprehensive genotyping of the USA national maize inbred seed bank. Genome Biol 14 10.1186/gb-2013-14-6-r55

Saglam A, Kadioglu A, Demiralay M, Terzi R (2014) Leaf rolling reduces photosynthetic loss in maize under severe drought. Acta Bot Croat 73:315–332 10.2478/botcro-2014-0012

Saitou N, Nei M (1987) The Neighbor-Joining Method - a New Method for Reconstructing Phylogenetic Trees. Mol Biol Evol 4:406–425 10.1093/oxfordjournals.molbev.a040454

Scheben A, Batley J, Edwards D (2017) Genotyping-by-sequencing approaches to characterize crop genomes: choosing the right tool for the right application. Plant Biotechnol J 15:149–161 10.1111/pbi.12645

Scott MF, Ladejobi O, Amer S, Bentley AR, Biernaskie J, Boden SA, Clark M, Dell’Acqua M, Dixon LE, Filippi CV, Fradgley N, Gardner KA, Mackay IJ, O’Sullivan D, Percival-Alwyn L, Roorkiwal M, Singh RK, Thudi M, Varshney RK, Venturini L, Whan A, Cockram J, Mott R (2020) Multi-parent populations in crops: a toolbox integrating genomics and genetic mapping with breeding. Heredity 125:396–416 10.1038/s41437-020-0336-6

Sekhon RS, Saski C, Kumar R, Flinn BS, Luo F, Beissinger TM, Ackerman AJ, Breitzman MW, Bridges WC, de Leon N, Kaepplere SM (2019) Integrated Genome-Scale Analysis Identifies Novel Genes and Networks Underlying Senescence in Maize. Plant Cell 31:1968–1989 10.1105/tpc.18.00930

Shah R, Huang BE, Whan A, Newberry M, Verbyla K, Morell MK, Cavanagh CR (2019) The complex genetic architecture of recombination and structural variation in wheat uncovered using a large 8-founder MAGIC population. bioRxiv:594317 10.1101/594317

Shen W, Le S, Li Y, Hu FQ (2016) SeqKit: A Cross-Platform and Ultrafast Toolkit for FASTA/Q File Manipulation. Plos One 11 10.1371/journal.pone.0163962

Song L, Florea L, Langmead B (2014) Lighter: fast and memory-efficient sequencing error correction without counting. Genome Biol 15 10.1186/s13059-014-0509-9

Tian F, Bradbury PJ, Brown PJ, Hung H, Sun Q, Flint-Garcia S, Rocheford TR, McMullen MD, Holland JB, Buckler ES (2011) Genome-wide association study of leaf architecture in the maize nested association mapping population. Nat Genet 43:159–U113 10.1038/ng.746

Tollenaar M, Wu J (1999) Yield improvement in temperate maize is attributable to greater stress tolerance. Crop science 39:1597–1604 10.2135/cropsci1999.3961597x

Unterseer S, Bauer E, Haberer G, Seidel M, Knaak C, Ouzunova M, Meitinger T, Strom TM, Fries R, Pausch H, Bertani C, Davassi A, Mayer KFX, Schön CC (2014) A powerful tool for genome analysis in maize: development and evaluation of the high density 600 k SNP genotyping array. Bmc Genomics 15 10.1186/1471-2164-15-823

VanRaden PM (2008) Efficient Methods to Compute Genomic Predictions. J Dairy Sci 91:4414–4423 10.3168/jds.2007-0980

Vogt F, Shirsekar G, Weigel D (2022) vcf2gwas: Python API for comprehensive GWAS analysis using GEMMA. Bioinformatics 38:839–840 10.1093/bioinformatics/btab710

Wald A (1943) Tests of statistical hypotheses concerning several parameters when the number of observations is large. Transactions of the American Mathematical society 54:426–482 10.1090/S0002-9947-1943-0012401-3

Wang SB, Xie FJ, Xu SZ (2022) Estimating genetic variance contributed by a quantitative trait locus: A random model approach. Plos Comput Biol 18 10.1371/journal.pcbi.1009923

Wright S (1978) Evolution and the genetics of populations. A treatise in four volumes. Volume 4. Variability within and among natural populations

Yang J, Zaitlen NA, Goddard ME, Visscher PM, Price AL (2014) Advantages and pitfalls in the application of mixed-model association methods. Nat Genet 46:100–106 10.1038/ng.2876

Zhang C, Dong SS, Xu JY, He WM, Yang TL (2019) PopLDdecay: a fast and effective tool for linkage disequilibrium decay analysis based on variant call format files. Bioinformatics 35:1786–1788 10.1093/bioinformatics/bty875

Zhou X, Stephens M (2012) Genome-wide efficient mixed-model analysis for association studies. Nat Genet 44:821–U136 10.1038/ng.2310

