## SupplementaryMethodFigureTable for "Genetic Dissection of Grain Yield and Correlated Proxy Traits Under Suboptimal Conditions"

### Supplementary Method

#### Method S1. IBD-based QTL mapping

For IBD-based QTL mapping, a subset of 19,981 (~20K) markers evenly covering the entire genome was analyzed to achieve a better balance between resolution and computational efficiency. Marker positions were projected onto a genetic map constructed from an $F_2$ population generated from a cross between flint and dent maize lines (EP1 × PH207) (Haberer et al., 2020).

QTL detection for individual trials was performed by the R package statgenMPP (Li et al., 2022) using founder (FH) and ancestral haplotype (AH) probabilities as genetic predictors. The model is expressed as follows:

$$\boldsymbol{y} = \mathbf{1}\mu + \mathbf{M_q}\boldsymbol{a_q} + \sum_{c \in \boldsymbol{C}} \mathbf{M_c}\boldsymbol{a_c} + \mathbf{Z}\boldsymbol{u} + \boldsymbol{\varepsilon} \qquad \text{(S1)}$$

where $\boldsymbol{y}$ is the $n$-dimensional vector of adjusted entry means, with $n$ being the number of MAGIC lines ($n$ = 388); $\mathbf{1}$ is a vector of ones, $\mu$ is the overall mean; $\boldsymbol{a_q}$ is a $p$-dimensional vector of random effects at the putative QTL $q$, with $p$ equal to the number of alleles (8 for founder alleles or ≤ 8 for ancestral alleles) with $\boldsymbol{a_q} \sim N(\mathbf{0}, \mathbf{I_p}\sigma_q^2)$; $\mathbf{M_q}$ is the $n$ x $p$ design matrix allocating the expected genotype score, calculated as twice the probability of a founder or ancestral allele at a putative QTL position; $\boldsymbol{a_c}$ is a $p$-dimensional vector of effects for a given cofactor c (representing other QTL), where $\boldsymbol{a_c} \sim N(\mathbf{0}, \mathbf{I_p}\sigma_c^2)$; $\mathbf{M_c}$ denotes the $n$ x $p$ design matrix corresponding to cofactor c; $\boldsymbol{u}$ is the $n$-dimensional vector of random polygenic effects, with $\boldsymbol{u} \sim N(0, \mathbf{K}\sigma_g^2)$; $\mathbf{Z}$ is the design matrix for the polygenic effects; and $\boldsymbol{e}$ is the $n$-dimensional vector of random residual effects, with $\boldsymbol{\varepsilon} \sim N(0, \mathbf{I_n}\sigma^{\mathbf{2}})$. $\mathbf{K}$, $\sigma_g^2$ and $\sigma^{\mathbf{2}}$ are defined analogously to Eq. (2).

We tested if the variance of the QTL ($\sigma_q^2$) differed significantly from zero with a likelihood ratio test (LRT) comparing the alternative ($\boldsymbol{a_q} \neq \mathbf{0}$) and null ($\boldsymbol{a_q} = \mathbf{0}$) model. A significance threshold of $-\log_{10}(p\text{-value}) = 4$ was inferred from 1,000 permutations using the FH method on two quantitative traits, testcross grain yield and line *per se* plant height. For both traits, similar thresholds of 3.8 (α = 0.05) and 4.7 (α = 0.01) were found and we consequently applied also to all the traits. The final QTL model was obtained through iterative stepwise forward selection, continuing until no additional QTL exceeded the significance threshold (Li et al., 2022). The peak markers on the logarithm of the odds (LOD) curves represented the genotypes of the detected QTL.

For obtaining final set QTL, we adapted the approach from Li et al. (2024) for the a multi-locus, multi-trial QTL model:

$$y_{ij} = \mu + t_j + \sum_{q \in \boldsymbol{Q}} \boldsymbol{m_q} \boldsymbol{a_{q(j)}} + u_i + \varepsilon_{ij} \quad \text{(S2)}$$

Where $y_{ij}$ is the adjusted mean of genotype $i$ in trial $j$; $\mu$ is the overall mean; $t_j$ is the fixed effect of trial $j$; $\boldsymbol{m_q}$ is a $p$-dimensional row vector of expected genotype scores, calculated as twice the IBD probabilities at a putative QTL position, where $p$ is the number of founder or ancestral alleles; $\boldsymbol{a_{q(j)}}$ is a $p$-dimensional column vector of random allelic effects at the putative QTL in trial $j$. For QTL with consistent effects across trial, allelic effects were assumed to be equal: $\boldsymbol{a_{q(1)}} = \boldsymbol{a_{q(2)}} = \cdots \boldsymbol{a_{q(j)}} = \boldsymbol{a_q}$, where $\boldsymbol{a_q} \sim N(\boldsymbol{0}, \mathbf{I_p}\sigma_q^2)$. For QTL with trial-specific effects, the allelic effects vary across trials: $\boldsymbol{a_q^j} \sim N(\boldsymbol{0}, \mathbf{I_p}\sigma_{q(j)}^2)$. The QTL effect type was determined by comparing the Akaike Information Criterion (AIC) of the two models and selecting the one with the lower AIC. $u_i$, $\mathbf{K}^*$, $\sigma_g^2$, $\varepsilon_{ij}$, $\sigma^2$, and $\boldsymbol{Q}^*$ are defined analogously to Eq. (4).

To avoid collinearity among QTL detected in different trials, peak markers of QTL located in close proximity (< 5 *cM*) were grouped together and the marker with the lowest *p*-value was retained for backward elimination. At each step, marker significance was assessed using likelihood ratio tests (LRTs), comparing models with and without the QTL effect. The marker with the highest *p*-value was removed iteratively until all remaining markers had *p*-values < 0.01. After backward elimination, we computed the LOD curves within a 10 *cM* window around each final QTL to refine peak positions and determine confidence intervals.

The proportion of phenotypic variance explained by QTL in individual trial analyses was estimated using Eq. (3), with variances components ($\sigma_q^2$, $\sigma_g^2$ and $\sigma^2$) defined analogously to those in Eq. (S1). Allelic effects were estimated as the best linear unbiased predictors (BLUPs) of $\boldsymbol{a_q}$ in Eq. (S1) and Eq. (S2), with effects constrained to sum to zero. Confidence intervals were defined using a 1.5 LOD-drop from the peak markers. The same QTL nomenclature as described for the SNP-based method was applied, with the additional suffix "-FH" or "-AH" appended.

**Method S2. Marker density comparison for QTL detection**

We generated two SNP datasets with different marker densities from the full 2.7 M SNP set: one overlapping with the 600k Affymetrix® Axiom® Maize Array (Unterseer et al., 2014) and the other with a 15K custom Illumina Array developed by KWS SAAT SE & Co. KGaA, resulting in 259,341 and 8,408 SNPs, respectively, for QTL analyses. Adjusted means of GDY across seven testcross trials were used as phenotypes. QTL detection followed the methods described in Eq. (2) for SNP- and Eq. (S1) for IBD-based methods.

#### Supplementary Table

**Table S1** Crossing scheme for the MAGIC population: Pedigree details of the three subfamilies and the corresponding number of DH lines derived from them (N).

| Subfamily | N |
|---|---|
| 1. PHG83/F888//Lo1056/Lo1270///FC1890/B106//B107/Lo1290 | 137 |
| 2. FC1890/B106//PHG83/F888///B107/Lo1290//Lo1056/Lo1270 | 154 |
| 3. FC1890/B106//Lo1056/Lo1270///B107/Lo1290//PHG83/F888 | 97 |
| **A**: PHG83, **B**: F888, **C**: Lo1056, **D**: Lo1270, **E**: FC1890, **F0**: B106, **G**: B107, **H**: Lo1290 | |

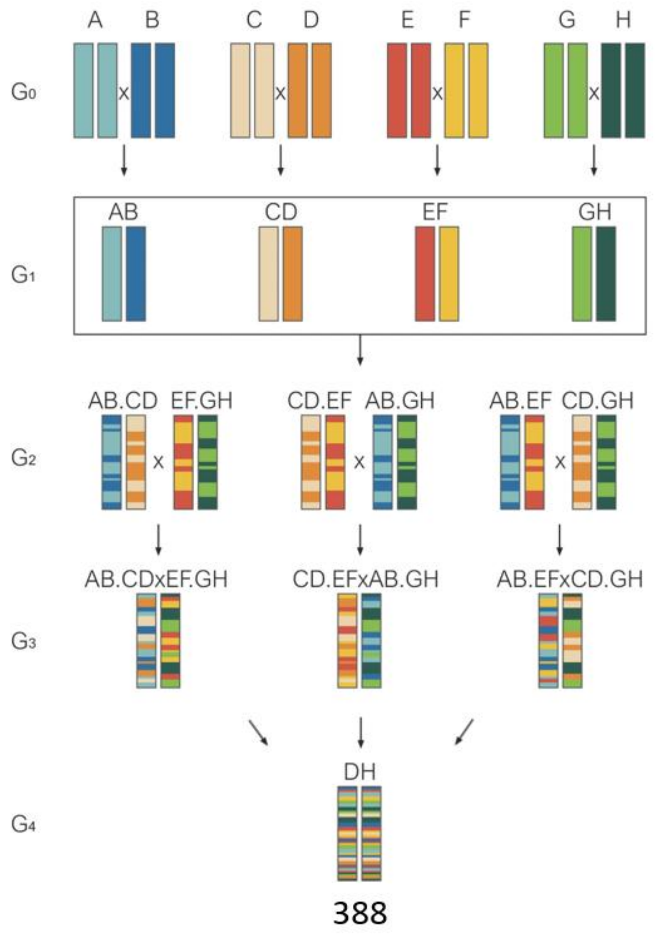

**Table S2** Overview of field trials. **Type:** testcross (TC) or line *per se* (LP). **Location:** KAB: Kaba, Hungary (47.36 ° N, 21.27° E); MUR: Murony, Hungary (46.77° N, 21.02° E); PNF: Pleinfeld, Germany (49.17° N, 11.04° E); DTH: Dietersheim, Germany (48.27° N, 11.04° E); MCE: Monselice, Italy (45.24° N, 11.75° E); MER: Merdingen, Germany (48.02° N, 7.69° E); ROG: Roggenstein, Germany (48.18° N, 11.32° E). **Management:** irrigated (I) or rainfed (R). **Plot:** long 2-rows plots for yield evaluation (Y), short 1-row (O) or 2-row (O*) plots for line *per se* evaluation. **Trait:** grain dry matter yield (GDY), leaf-senescence (LS), leaf-rolling (LR), anthesis-silking interval (ASI), final plant height (PH_final), ear height (EH), female flowering (FF), male flowering (MF), grain dry matter content (GDC) and early plant height (PH_V6).

| Year | Type | Location | Management | Plot | Trait |
|---|---|---|---|---|---|
| 2020 | TC | KAB | I | Y | EH, PH_final, MF, FF, ASI, GDC, GDY |
| | | | R | Y | EH, PH_final, MF, FF, ASI, GDC, GDY |
| | | MUR | R | Y | EH, PH_final, MF, FF, ASI, GDC, GDY, LS |
| 2021 | TC | MUR | I | Y | PH_V6, MF, FF, ASI, GDC, GDY |
| | | | R | Y | PH_V6, EH, PH_final, MF, FF, ASI, GDC, GDY, LR, LS |
| | | PNF | R | Y | PH_V6, EH, PH_final, MF, FF, ASI, GDC, GDY |
| | | DTH | R | Y | PH_V6, EH, PH_final, MF, FF, ASI, GDC, GDY |
| | LP | MCE | I | O | PH_V6, EH, PH_final, MF, FF, ASI |
| | | | R | O | PH_V6, EH, PH_final, MF, FF, ASI, LR |
| | | MUR | I | O | PH_V6, EH, PH_final, MF, FF, ASI |
| | | | R | O | PH_V6, EH, PH_final, MF, FF, ASI, LR, LS |
| 2022 | LP | MCE | I | O | EH, PH_final, MF, FF, ASI |
| | | | R | O | EH, PH_final, MF, FF, ASI, LS |
| | | MER | I | Y | MF, FF, ASI, GDC, GDY |
| | | | R | Y | MF, FF, ASI, LS |
| | | MUR | I | Y | EH, PH_final, MF, FF, ASI, GDC, GDY |
| | | | R | Y | EH, PH_final, MF, LR, LS |
| | | DTH | R | Y | PH_V6, PH_final, MF, FF, ASI, GDC, GDY, LR, LS |
| | | ROG | I | Y | PH_V6, PH_final, MF, FF, ASI, GDC, GDY |
| 2023 | LP | DTH | R | O* | PH_V6, EH, PH_final, MF, FF, ASI, LR, LS |
| | | ROG | I | O* | PH_V6, EH, PH_final, MF, FF, ASI |

**Table S3** Variance component estimation of traits. Table includes number of trials (**nTrial**), genetic ($\boldsymbol{\sigma_g^2}$) and genotype-by-trial ($\boldsymbol{\sigma_{gt}^2}$) variance components, including standard errors, and their ratio ($\boldsymbol{\sigma_g^2}$**:**$\boldsymbol{\sigma_{gt}^2}$), heritability ($\boldsymbol{h^2}$) across trials and their 95% confidence interval ($\boldsymbol{h^2}$ **CI**), range (minimum and maximum) of repeatability from individual trials (**Repeatability**). Traits are grain dry matter yield (GDY), grain dry matter content (GDC), leaf-senescence (LS), leaf-rolling (LR), anthesis-silking interval (ASI), female flowering (FF), male flowering (MF), final plant height (PH_final), ear height (EH) and early plant height (PH_V6).

| Type | Trait | nTrial | $\sigma_g^2 \pm$ SE | $\sigma_{gt}^2 \pm$ SE | $\sigma_{gt}^2 : \sigma_g^2$ | $h^2$ | $h^2$ CI | Repeatability |
|---|---|---|---|---|---|---|---|---|
| Testcross | GDY | 7 | 0.54 ± 0.05 | 0.55 ± 0.03 | 1.03 | 0.81 | 0.78 - 0.83 | 0.41 - 0.79 |
| | GDC | 7 | 0.47 ± 0.04 | 0.49 ± 0.02 | 1.04 | 0.84 | 0.82 - 0.86 | 0.64 - 0.90 |
| | LS | 2 | 0.14 ± 0.03 | 0.12 ± 0.02 | 0.88 | 0.52 | 0.41 - 0.60 | 0.36 - 0.54 |
| | LR | 1 | - | - | - | - | - | 0.45 |
| | ASI | 7 | 0.15 ± 0.02 | 0.11 ± 0.02 | 0.72 | 0.70 | 0.65 - 0.74 | 0.06 - 0.56 |
| | FF | 7 | 1.12 ± 0.09 | 0.31 ± 0.03 | 0.28 | 0.92 | 0.90 - 0.93 | 0.39 - 0.77 |
| | MF | 7 | 0.62 ± 0.05 | 0.18 ± 0.02 | 0.29 | 0.89 | 0.88 - 0.91 | 0.23 - 0.67 |
| | PH_final | 6 | 84.89 ± 6.88 | 15.78 ± 2.10 | 0.19 | 0.91 | 0.90 - 0.93 | 0.54 - 0.75 |
| | EH | 6 | 63.18 ± 5.18 | 10.33 ± 1.77 | 0.16 | 0.90 | 0.89 - 0.92 | 0.41 - 0.68 |
| | PH_V6 | 4 | 4.56 ± 0.55 | 2.33 ± 0.43 | 0.51 | 0.73 | 0.67 - 0.77 | 0.38 - 0.50 |
| Line *per se* | GDY | 4 | 0.49 ± 0.07 | 0.56 ± 0.05 | 1.14 | 0.70 | 0.64 - 0.75 | 0.18 - 0.84 |
| | GDC | 4 | 17.87 ± 2.03 | 11.37 ± 1.09 | 0.64 | 0.81 | 0.77 - 0.84 | 0.38 - 0.95 |
| | LS | 6 | 0.34 ± 0.03 | 0.35 ± 0.02 | 1.02 | 0.78 | 0.75 - 0.81 | 0.48 - 0.67 |
| | LR | 5 | 0.79 ± 0.08 | 0.68 ± 0.05 | 0.86 | 0.78 | 0.74 - 0.81 | 0.50 - 0.78 |
| | ASI | 13 | 2.18 ± 0.18 | 0.99 ± 0.06 | 0.46 | 0.92 | 0.91 - 0.93 | 0.34 - 0.72 |
| | FF | 13 | 9 ± 0.68 | 2.19 ± 0.10 | 0.24 | 0.97 | 0.96 - 0.97 | 0.57 - 0.91 |
| | MF | 14 | 6.52 ± 0.49 | 1.73 ± 0.07 | 0.27 | 0.97 | 0.96 - 0.97 | 0.46 - 0.88 |
| | PH_final | 12 | 289.04 ± 21.73 | 56.83 ± 2.84 | 0.20 | 0.97 | 0.97 - 0.97 | 0.54 - 0.89 |
| | EH | 10 | 143.85 ± 10.84 | 22.19 ± 1.51 | 0.15 | 0.97 | 0.96 - 0.97 | 0.57 - 0.84 |
| | PH_V6 | 8 | 14.07 ± 1.20 | 5.98 ± 0.57 | 0.42 | 0.87 | 0.85 - 0.89 | 0.34 - 0.69 |

**Table S4** Summary of QTL. **nTrial** refers to the number of tested trials. Columns **SNP**, **FH**, and **AH** indicate the number of QTL of final set identified by each method, and numbers in parentheses represent total number of QTL detected from individual trials, and $\boldsymbol{\sigma_g^2}$ **explained** indicates the proportion of genetic variance explained by all QTL identified using the SNP method. Traits are grain dry matter yield (GDY), grain dry matter content (GDC), leaf-senescence (LS), leaf-rolling (LR), anthesis-silking interval (ASI), female flowering (FF), male flowering (MF), final plant height (PH_final), ear height (EH) and early plant height (PH_V6). *Testcross leaf rolling was measured in a single trial; therefore, only individual-trial QTL counts are reported.

| | | nTrial | SNP | FH | AH | $\sigma_g^2$ explained |
|---|---|---|---|---|---|---|
| Testcross | GDY | 7 | 22 (71) | 10 (24) | 7 (22) | 0.45 |
| | GDC | 7 | 34 (147) | 11 (51) | 10 (51) | 0.53 |
| | LS | 2 | 13 (20) | 5 (6) | 5 (6) | 0.45 |
| | *LR | 1 | - (5) | - (2) | - (2) | - (0.50) |
| | ASI | 7 | 9 (13) | 2 (3) | 2 (4) | 0.53 |
| | FF | 7 | 41 (122) | 10 (40) | 13 (38) | 0.34 |
| | MF | 7 | 32 (132) | 9 (38) | 10 (43) | 0.43 |
| | PH_final | 6 | 21 (64) | 5 (19) | 7 (26) | 0.55 |
| | EH | 6 | 25 (96) | 8 (24) | 11 (32) | 0.52 |
| | PH_V6 | 4 | 0 (0) | 1 (2) | 1 (3) | 0 |
| Line *per se* | GDY | 4 | 0 (0) | 1 (1) | 4 (6) | 0 |
| | GDC | 4 | 17 (31) | 4 (7) | 5 (10) | 0.28 |
| | LS | 6 | 24 (36) | 5 (9) | 4 (7) | 0.53 |
| | LR | 5 | 22 (47) | 7 (17) | 7 (16) | 0.3 |
| | ASI | 13 | 15 (77) | 3 (26) | 1 (27) | 0.7 |
| | FF | 13 | 30 (190) | 7 (58) | 7 (56) | 0.66 |
| | MF | 14 | 29 (202) | 5 (38) | 2 (39) | 0.75 |
| | PH_final | 12 | 11 (49) | 4 (40) | 5 (39) | 0.68 |
| | EH | 10 | 23 (153) | 4 (34) | 4 (36) | 0.58 |
| | PH_V6 | 8 | 14 (41) | 4 (13) | 4 (14) | 0.6 |

**Table S5** Prediction accuracy of single-trait GBLUP using genomic relationship matrices derived from **SNP**, founder and ancestral haplotype probabilities (**FH**, and **AH**) using adjusted means across trials. Values represent means with standard deviations from 100 cross validation iterations. Traits are grain dry matter yield (GDY), grain dry matter content (GDC), leaf-senescence (LS), leaf-rolling (LR), anthesis-silking interval (ASI), female flowering (FF), male flowering (MF), final plant height (PH_final), ear height (EH) and early plant height (PH_V6).

| | | **SNP** | **FH** | **AH** |
|---|---|---|---|---|
| Testcross | GDY | 0.60 ± 0.05 | 0.60 ± 0.05 | 0.61 ± 0.05 |
| | GDC | 0.74 ± 0.04 | 0.72 ± 0.04 | 0.73 ± 0.04 |
| | ASI | 0.52 ± 0.07 | 0.48 ± 0.07 | 0.51 ± 0.07 |
| | LS | 0.33 ± 0.09 | 0.34 ± 0.09 | 0.32 ± 0.08 |
| | FF | 0.67 ± 0.05 | 0.63 ± 0.05 | 0.66 ± 0.05 |
| | MF | 0.69 ± 0.04 | 0.66 ± 0.04 | 0.69 ± 0.04 |
| | PH_final | 0.57 ± 0.05 | 0.53 ± 0.05 | 0.57 ± 0.05 |
| | EH | 0.62 ± 0.04 | 0.60 ± 0.05 | 0.62 ± 0.04 |
| | PH_V6 | 0.45 ± 0.12 | 0.46 ± 0.13 | 0.46 ± 0.13 |
| Line *per se* | GDY | 0.44 ± 0.09 | 0.43 ± 0.09 | 0.44 ± 0.09 |
| | GDC | 0.47 ± 0.11 | 0.45 ± 0.11 | 0.46 ± 0.11 |
| | LS | 0.40 ± 0.06 | 0.38 ± 0.06 | 0.40 ± 0.06 |
| | LR | 0.60 ± 0.05 | 0.61 ± 0.05 | 0.60 ± 0.05 |
| | ASI | 0.52 ± 0.05 | 0.50 ± 0.05 | 0.51 ± 0.05 |
| | FF | 0.61 ± 0.04 | 0.59 ± 0.05 | 0.61 ± 0.04 |
| | MF | 0.56 ± 0.05 | 0.55 ± 0.05 | 0.56 ± 0.05 |
| | PH_final | 0.56 ± 0.05 | 0.55 ± 0.05 | 0.56 ± 0.05 |
| | EH | 0.59 ± 0.05 | 0.58 ± 0.05 | 0.59 ± 0.05 |
| | PH_V6 | 0.51 ± 0.05 | 0.50 ± 0.05 | 0.50 ± 0.06 |

**Table S6** Co-localized grain yield QTL from the DROPS panel (Millet et al., 2016) and the MAGIC population. **QTL (DROPS)** refers to SNP marker of the QTL detected in the DROPS panel, and **CHR** and **POS** indicate their chromosome and physical position on the B73v5 reference genome. **QTL (MAGIC)** lists all testcross grain yield QTL found in MAGIC overlapping with the corresponding DROPS QTL.

| QTL (DROPS) | CHR | POS | QTL (MAGIC) |
|---|---|---|---|
| AX-90832809 | chr3 | 152503345 | *qGDY(TC)03B-FH* |
| AX-91388323 | chr3 | 17408785 | *qGDY(TC)03A-FH; qGDY(TC)03A-AH* |
| AX-90539808 | chr3 | 192515085 | *qGDY(TC)03C-FH* |
| AX-90973370 | chr5 | 209361458 | *qGDY(TC)05B* |
| AX-90548584 | chr6 | 25135854 | *qGDY(TC)06A; qGDY(TC)06B; qGDY(TC)06A-FH; qGDY(TC)06A-AH* |
| AX-91685880 | chr6 | 33061089 | *qGDY(TC)06A; qGDY(TC)06B* |
| AX-91685911 | chr6 | 33369214 | *qGDY(TC)06A; qGDY(TC)06B* |

#### Supplementary Figure

A

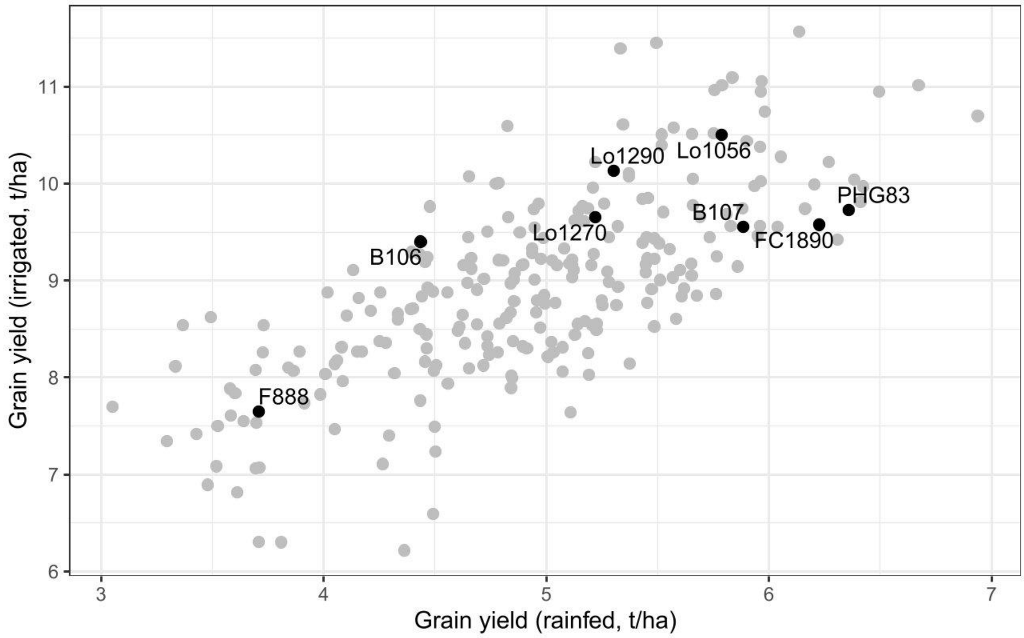


B

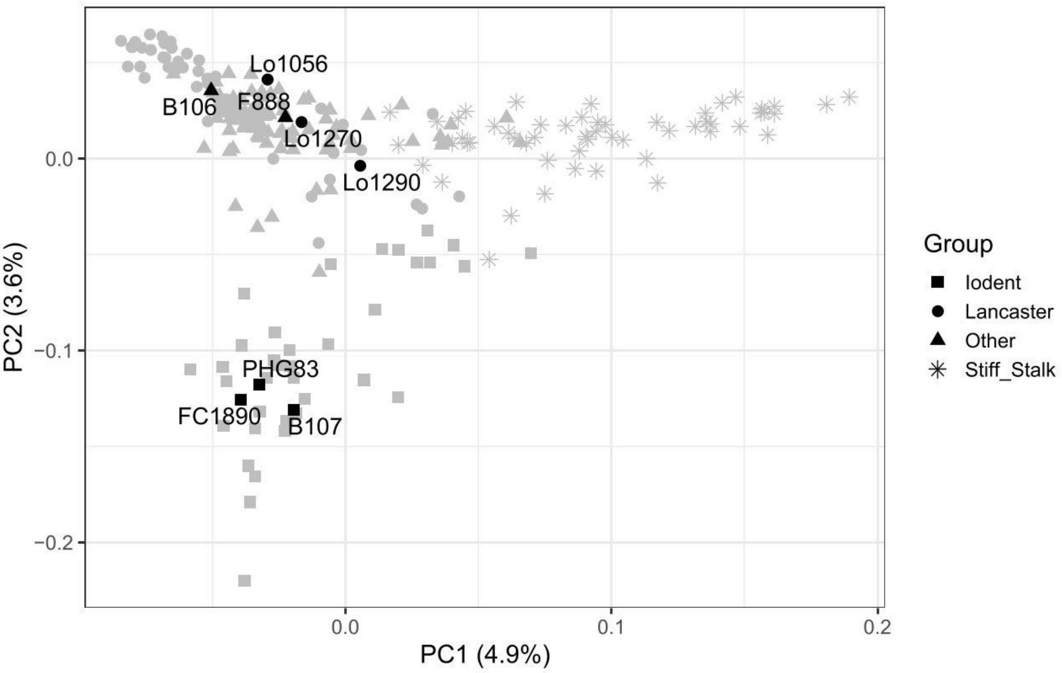


**Fig. S1** Selection of MAGIC founders. (A) Grain yield (t/ha) performance of the dent diversity panel (DROPS) with the eight MAGIC founders highlighted. The x-axis shows mean performance in rainfed trials across five environments (year-location combinations), and the y-axis shows mean performance in irrigated trials of the same environments. (B) First two principal coordinates of principal coordinate analysis (PCoA) based on modified Rogers' distance, calculated from 600K SNP array genotypic data. MAGIC founders are highlighted. Phenotypic and genotypic data from Millet et al. (2016).

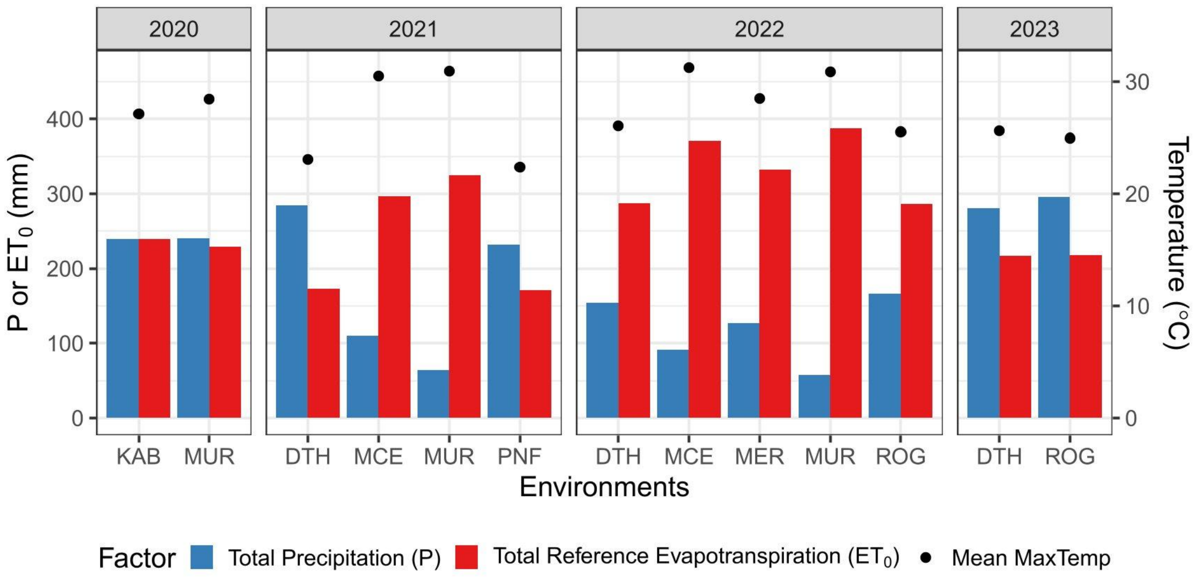


**Fig. S2** Weather conditions of all 13 year-location combinations during the 60-day period before and after male flowering (median across all plots). Bars show total precipitation (P) and reference evapotranspiration ($ET_0$) on the left y-axis, while dots indicate the average of the daily maximum temperature on the right y-axis.

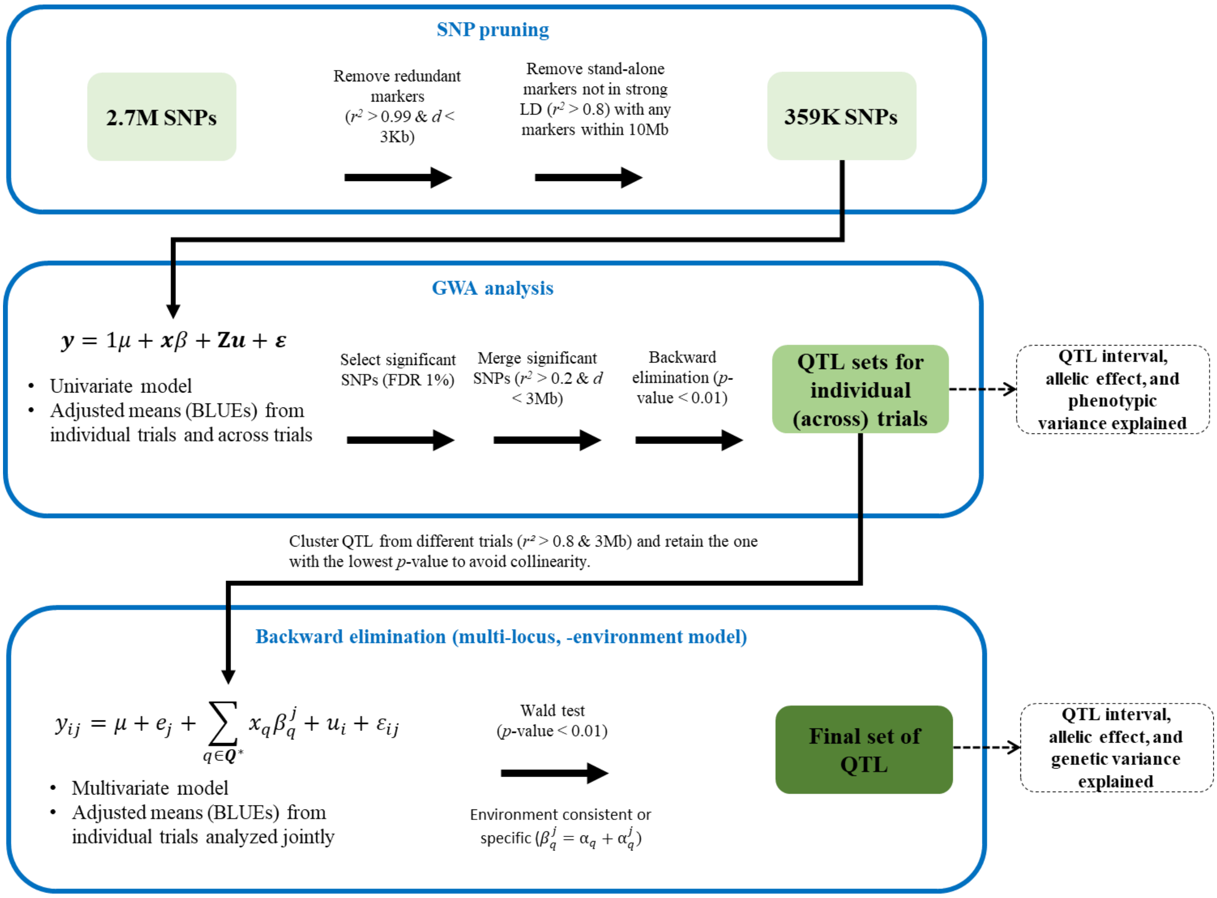


**Fig. S3** Flowchart of SNP-based QTL analyses. (A) Linkage disequilibrium (LD) pruning to remove redundant markers and filtering of stand-alone SNPs. In total, 359 k of 2.7 M SNPs remained for QTL analyses. (B) Genome-wide association (GWA) analyses conducted with adjusted means from individual trials as well as for across trials. Significant SNPs were merged, followed by backward elimination to obtain a set of QTL (SNPs) for each trait within each trial. (C) Multi-locus, multi-trial model implementing backward elimination to define the final set of QTL. Significance testing of QTL-by-trial (QxT) interactions was performed to determine whether QTLs were trial-consistent or trial-specific. See Methods for model notations.

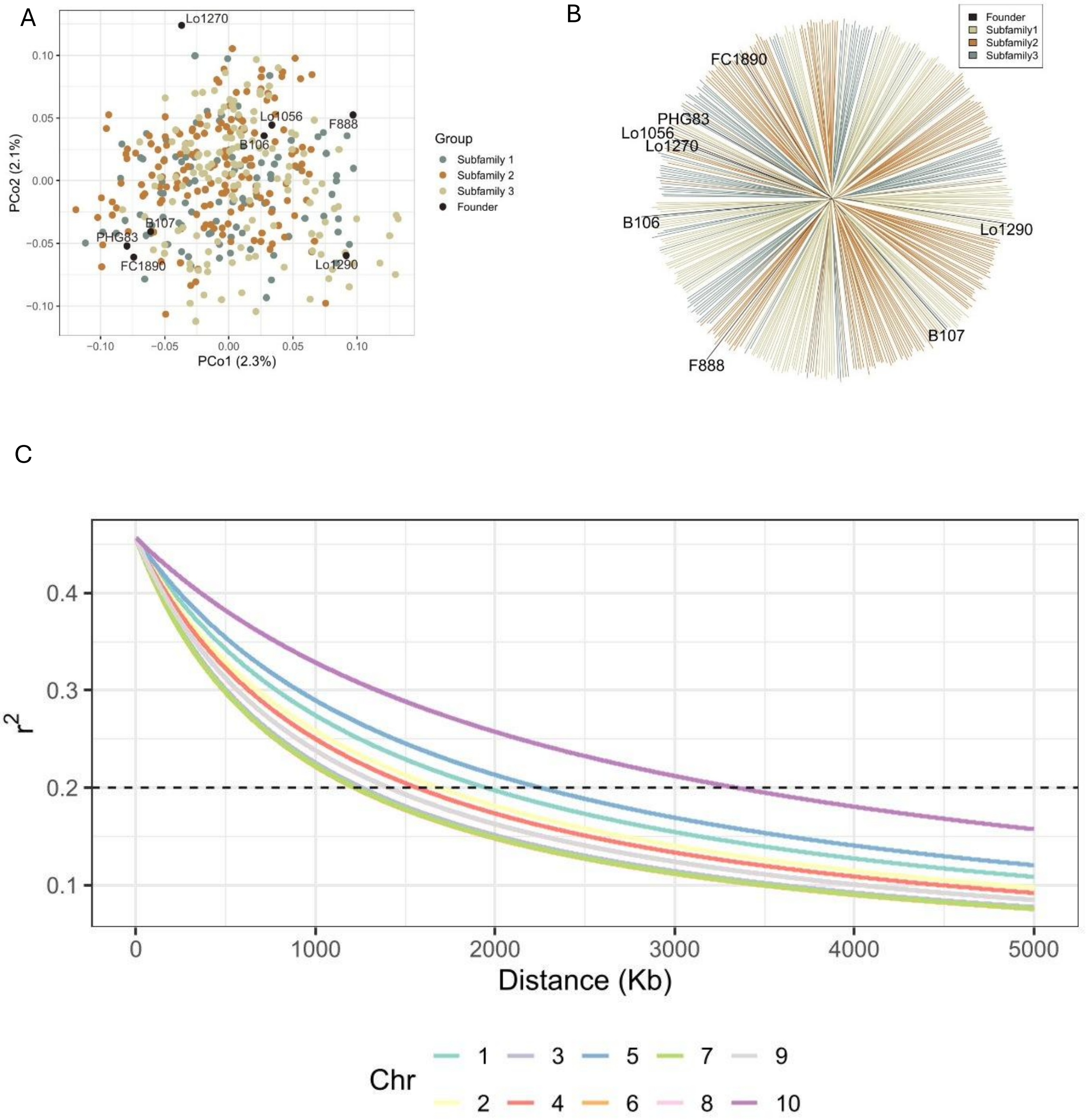


**Fig. S4** Molecular representation of the eight founders and the MAGIC 388 DH lines. Color indicates either the three subfamilies (Table S1) or the founders. (A) Principal coordinate analysis based on modified Rogers' distance, the first two principal coordinates are shown, with each genotype represented by a dot. (B) Phylogenetic tree comprising the founders and the DH lines (C) Linkage disequilibrium decay of the ten maize chromosomes represented by different colors.

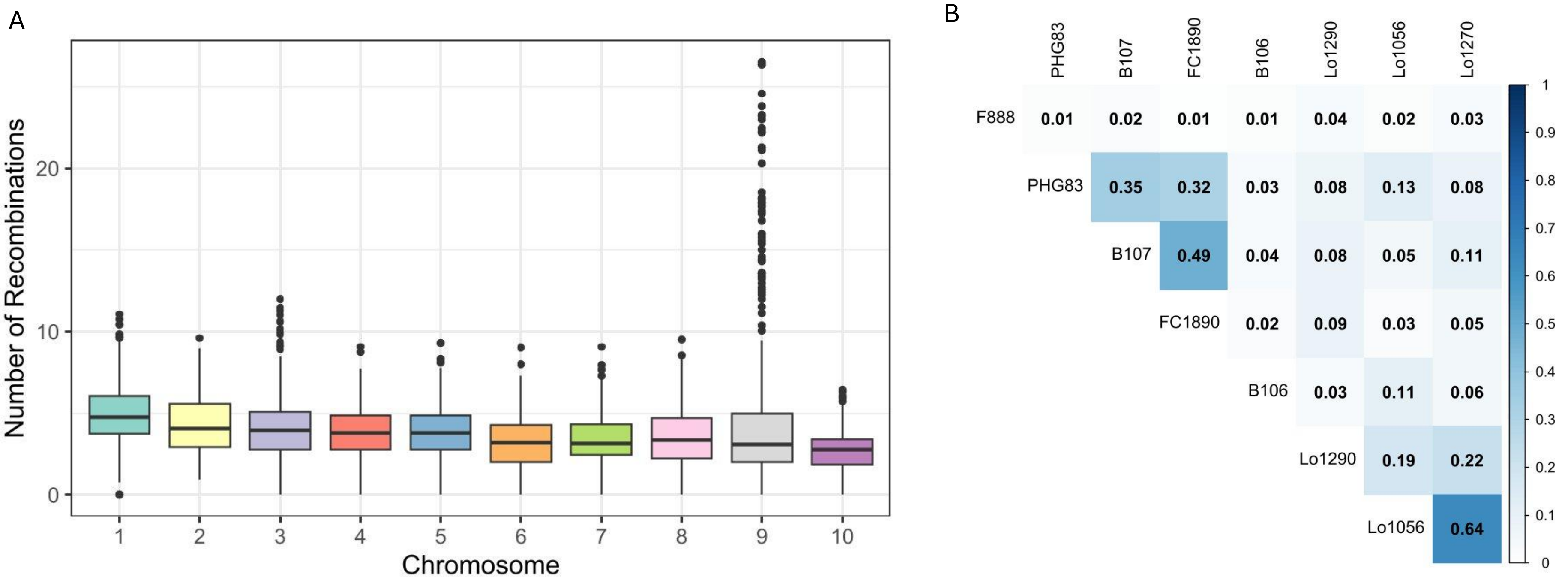


**Fig. S5** (A) Number of detected recombination events in the MAGIC 388 DH lines shown individually for the ten maize chromosomes. (B) Proportion of the genome identical by descent (IBD) for the 28 possible pairs of MAGIC founder lines.

GDY

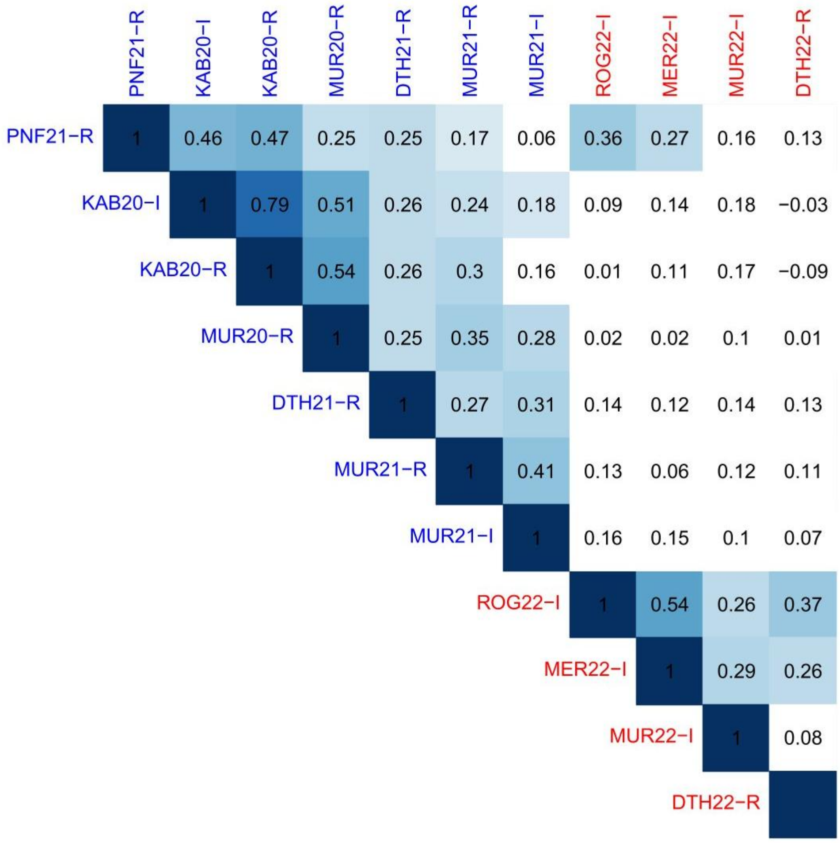


**Fig S6A** Phenotypic correlation heatmaps of grain dry matter yield (GDY) across all trials. Testcross trials are shown in blue, and line *per se* trials in red. Number indicates Pearson correlation coefficients, and colored grids denote significant correlations ($p < 0.01$) after multiple testing correction. Trials are ordered based on the loadings of the first principal component (**FPC** ordering method by R package *corrplot*)

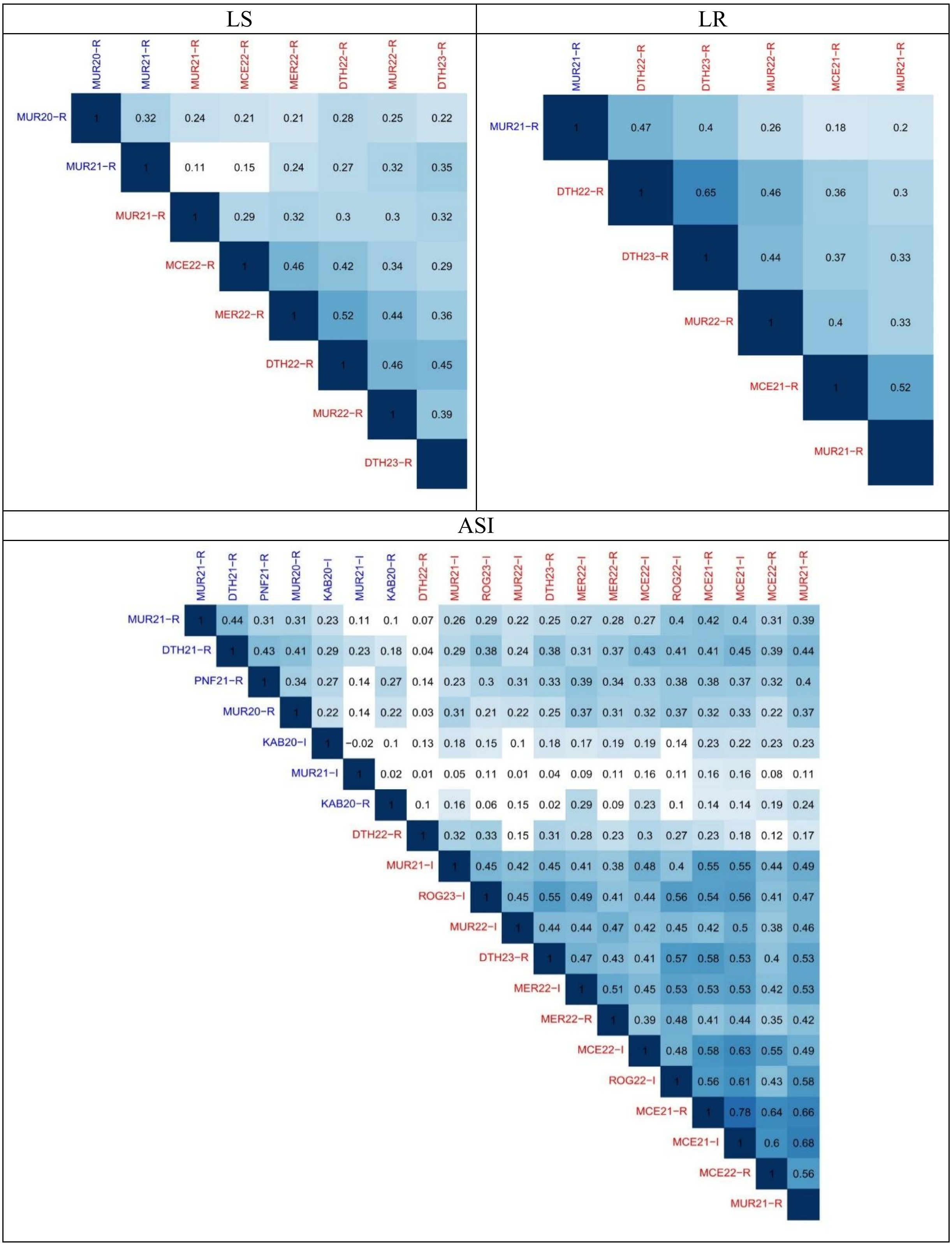


**Fig S6B** Phenotypic correlation heatmaps of leaf senescence (LS), leaf rolling (LR) and anthesis-silking interval (ASI) across all trials. Testcross trials are shown in blue, and line *per se* trials in red. Number indicates Pearson correlation coefficients, and colored grids denote significant correlations ($p < 0.01$) after multiple testing correction. Trials are ordered based on the loadings of the first principal component (**FPC** ordering method by R package *corrplot*).

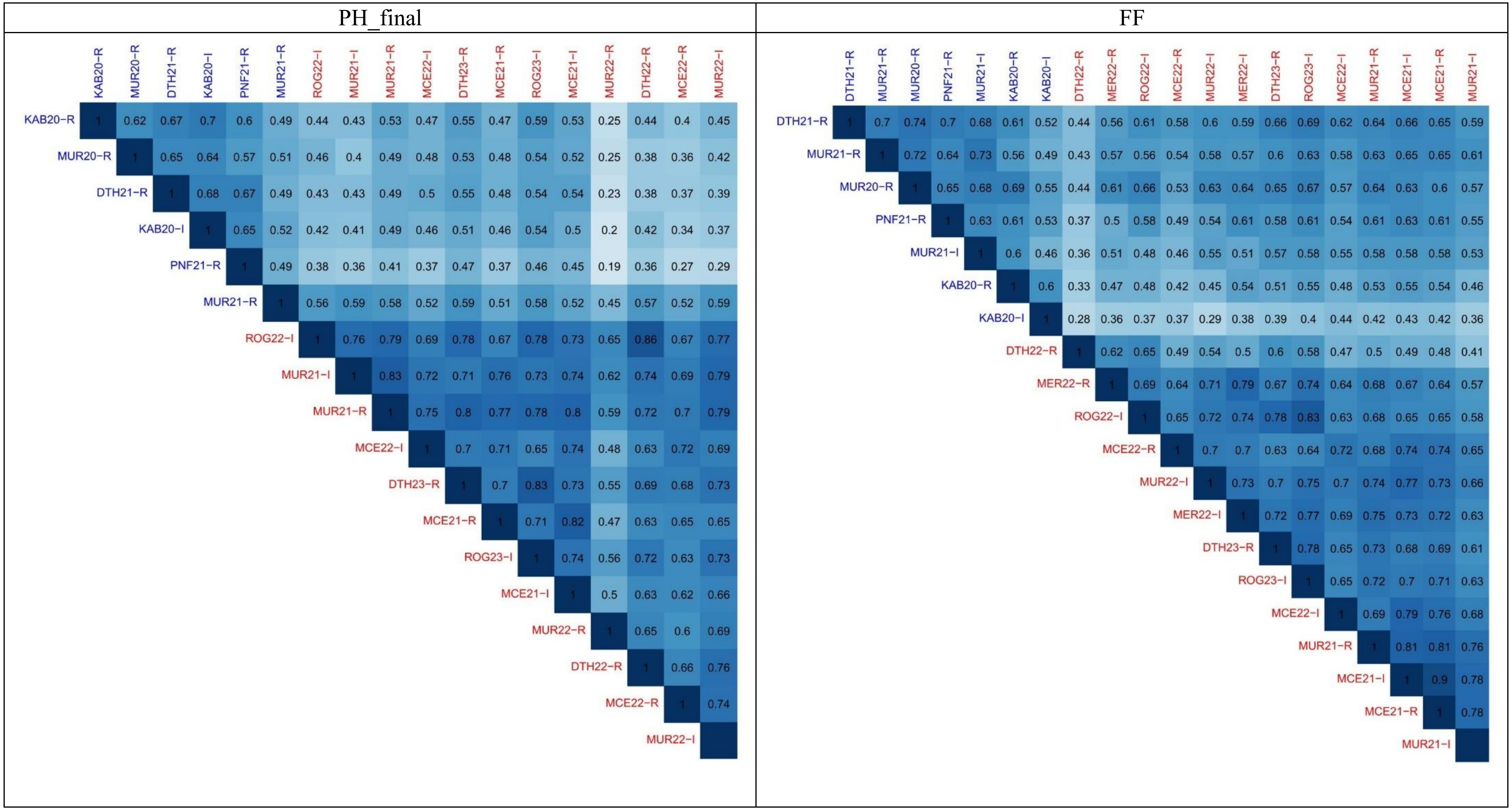


**Fig S6C** Phenotypic correlation heatmaps of final plant height (PH_final) and female flowering time (FF) across all trials. Testcross trials are shown in blue, and line *per se* trials in red. Number indicates Pearson correlation coefficients, and colored grids denote significant correlations ($p < 0.01$) after multiple testing correction. Trials are ordered based on the loadings of the first principal component (**FPC** ordering method by R package *corrplot*).

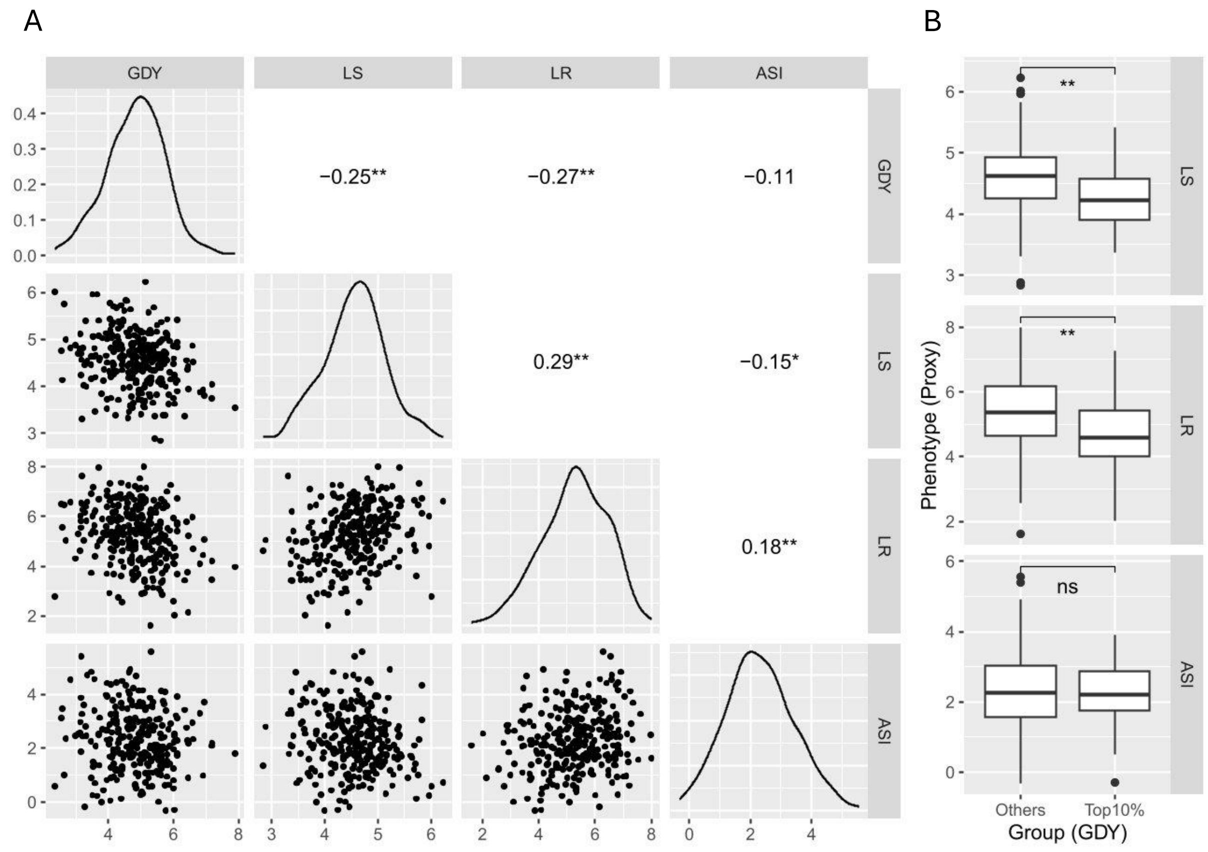


**Fig. S7** (A) Pairwise phenotypic correlations among grain dry matter yield (GDY), leaf senescence (LS), leaf rolling (LR) and anthesis-silking interval (ASI) in testcrosses evaluated in Murany (2021) under rainfed conditions. The upper triangle shows Pearson correlation coefficients. Diagonal panels display trait distributions. The lower triangle panels show pairwise scatter plots. (B) Boxplots comparing proxy traits performance between the top 10% highest-yielding genotypes and the remaining genotypes in the population. Asterisks indicate significance based on t-tests: $p > 0.05$ (ns), $p < 0.05$ (*), $p < 0.01$ (**).

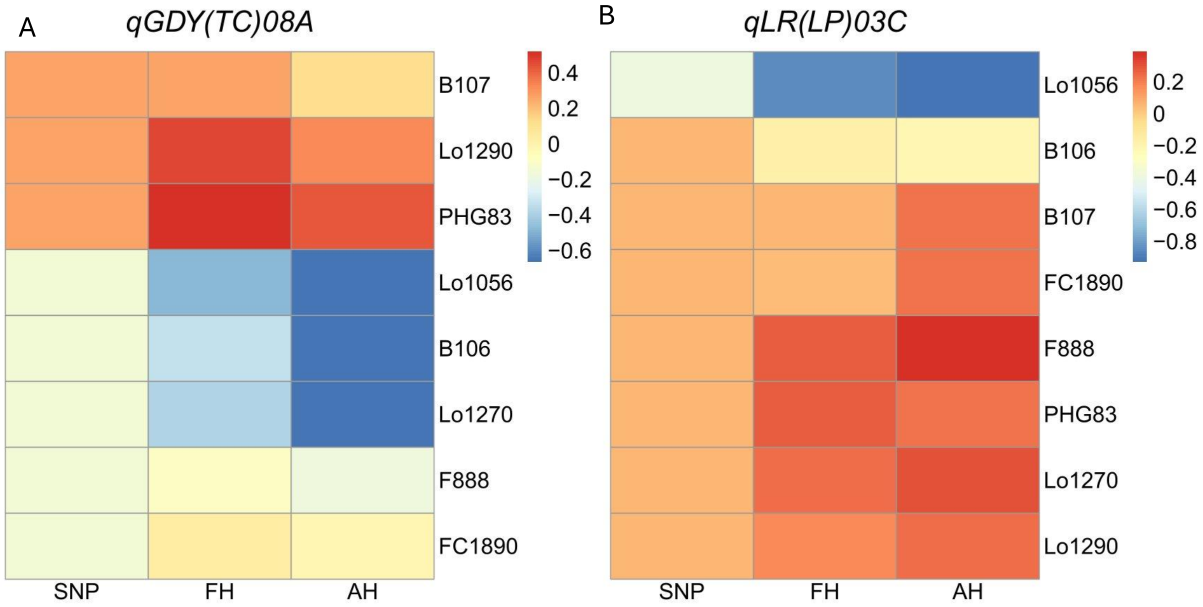


**Fig. S8** Heatmaps of allelic effect estimates for (A) a grain yield QTL, *qGDY(TC)08A*, affecting testcross grain yield and (B) a leaf rolling QTL, *qLR(LP)03C*, affecting line *per se* leaf rolling performance. Both QTL were detected all three mapping methods (SNP, FH and AH) and classified as QTL with consistent effect across trials. Columns represent different QTL detection methods, and rows correspond to effect estimates for alleles of the eight founders. Units are given on the original scale.

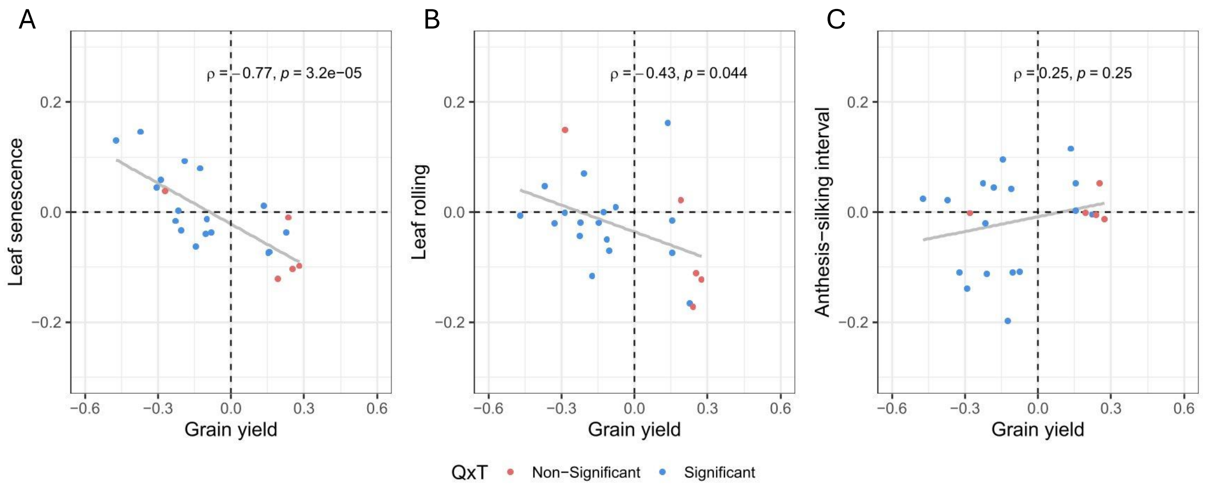


**Fig. S9** Correlation of SNP effects between grain yield and proxy traits at genomic positions of lead SNPs identified for grain yield QTL. The x-axis shows SNP effect estimates on grain yield across all testcross trials, while the y-axis shows effects of the same SNP on (A) leaf senescence and (B) leaf rolling and (C) anthesis-silking interval across all line *per se* trials. Pearson correlation coefficients and associated significance levels are shown in each panel; Dot colors indicate whether the QTL exhibits significant QTL-by-Trial (QxT) interaction.

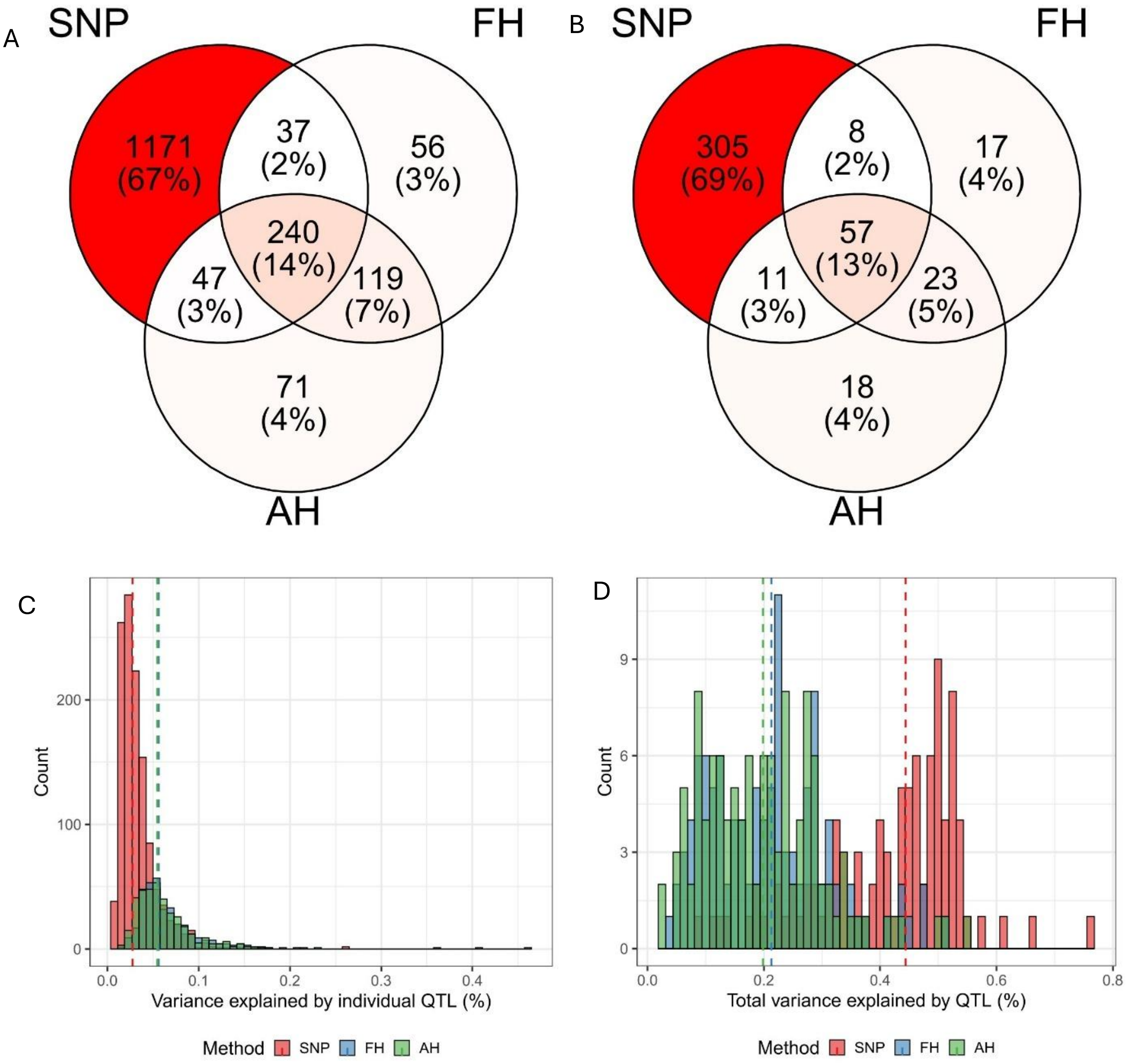


**Fig. S10** Comparison of QTL detection across three methods based on SNPs, founder (FH) or ancestral (AH) haplotypes. QTL overlap is shown by Venn diagrams: (A) single-trial analyses (140 trait-trial combinations) and (B) multi-trial analyses (19 traits). QTL of same trait with overlapping confidence intervals are considered identical. Panels (C) and (D) show the distribution of the proportion of phenotypic variance explained by QTL in single-trial analyses for individual QTL (C) and all QTL fitted simultaneously in the model (D). Dashed vertical lines mark the median proportion of variance explained for each method.

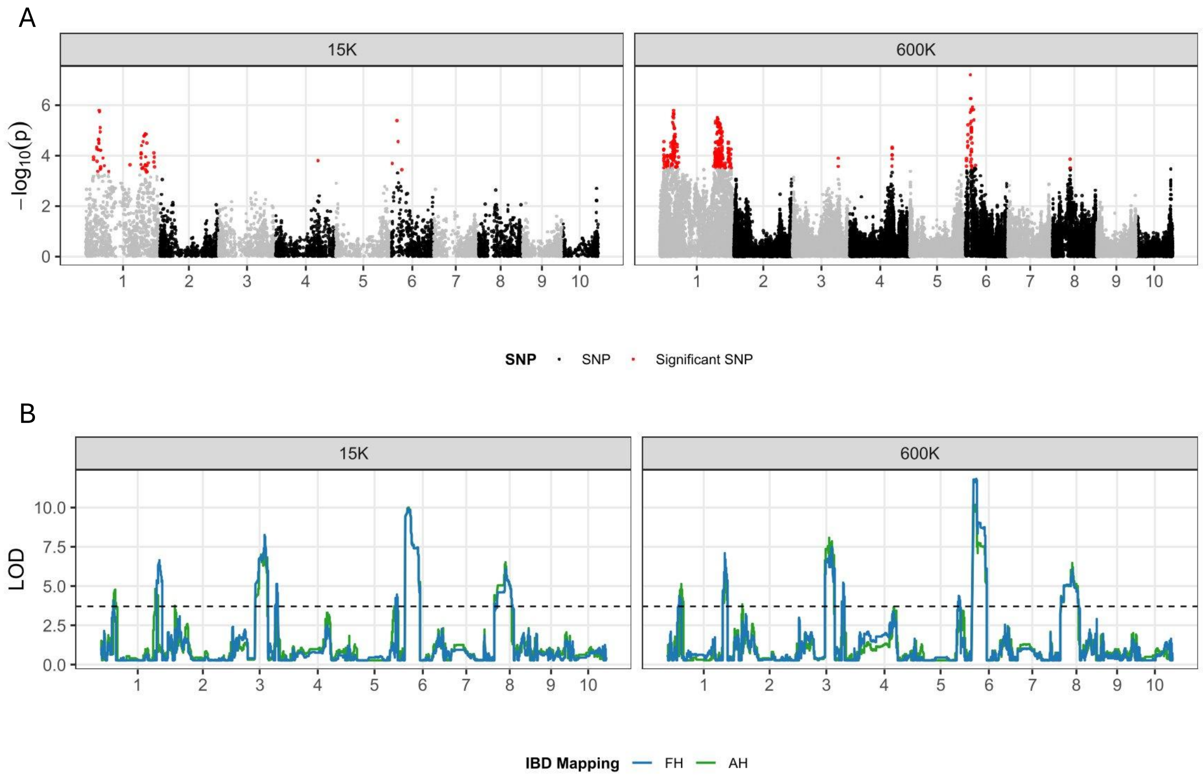


**Fig. S11** Comparison of QTL detection for grain yield across trials between 15k and 600k marker densities. (A) GWAS Manhattan plots, with significant SNPs (FDR < 0.01) shown in red. (B) IBD-based QTL mapping (AH and FH methods), with a significance threshold of $p < 0.0001$.

Haberer, G., Kamal, N., Bauer, E., Gundlach, H., Fischer, I., Seidel, M. A., Spannagl, M., Marcon, C., Ruban, A., Urbany, C., Nemri, A., Hochholdinger, F., Ouzunova, M., Houben, A., Schön, C. C., & Mayer, K. F. X. (2020). European maize genomes highlight intraspecies variation in repeat and gene content. *Nature Genetics*, *52*(9), 950–+. https://doi.org/10.1038/s41588-020-0671-9

Li, W. H., Boer, M. P., Joosen, R. V. L., Zheng, C. Z., Percival-Alwyn, L., Cockram, J., & Van Eeuwijk, F. A. (2024). Modeling QTL-by-environment interactions for multi-parent populations. *Frontiers in Plant Science*, *15*, Article 1410851. https://doi.org/10.3389/fpls.2024.1410851

Li, W. H., Boer, M. P., van Rossum, B. J., Zheng, C. Z., Joosen, R. V. L., & van Eeuwijk, F. A. (2022). statgenMPP: an R package implementing an IBD-based mixed model approach for QTL mapping in a wide range of multi-parent populations. *Bioinformatics*, *38*(22), 5134–5136. https://doi.org/10.1093/bioinformatics/btac662

Millet, E. J., Welcker, C., Kruijer, W., Negro, S., Coupel-Ledru, A., Nicolas, S. D., Laborde, J., Bauland, C., Praud, S., Ranc, N., Presterl, T., Tuberosa, R., Bedo, Z., Draye, X., Usadel, B., Charcosset, A., Van Eeuwijk, F., & Tardieu, F. (2016). Genome-Wide Analysis of Yield in Europe: Allelic Effects Vary with Drought and Heat Scenarios. *Plant Physiology*, *172*(2), 749–764. https://doi.org/10.1104/pp.16.00621

Unterseer, S., Bauer, E., Haberer, G., Seidel, M., Knaak, C., Ouzunova, M., Meitinger, T., Strom, T. M., Fries, R., Pausch, H., Bertani, C., Davassi, A., Mayer, K. F. X., & Schön, C. C. (2014). A powerful tool for genome analysis in maize: development and evaluation of the high density 600 k SNP genotyping array. *Bmc Genomics*, *15*, Article 823. https://doi.org/10.1186/1471-2164-15-823
